# NHJ-1 regulates canonical non-homologous end joining in *Caenorhabditis elegans*

**DOI:** 10.1101/763235

**Authors:** Aleksandar Vujin, Steven J. Jones, Monique Zetka

**Affiliations:** Department of Biology, McGill University, Montreal, QC, Canada; Canada’s Michael Smith Genome Sciences Centre, BC Cancer Agency, Vancouver, BC, Canada

## Abstract

Canonical non-homologous end joining (cNHEJ) is a near-universally conserved pathway for the repair of DNA double-strand breaks (DSBs). While the cNHEJ pathway encompasses more than a dozen factors in vertebrates and is similarly complex in other eukaryotes, in the nematode *C. elegans* the entire known cNHEJ toolkit consists of two proteins that comprise the Ku ring complex, *cku-70* and *cku-80*, and the terminal ligase *lig-4*. Here, we report the discovery of *nhj-1* as the fourth cNHEJ factor in *C. elegans*. Observing a difference in the phenotypic response to ionizing radiation (IR) between two lines of the wild type N2 strain, we mapped the locus causative of IR-sensitivity to a candidate on chromosome V. Using CRISPR-Cas9 mutagenesis, we show that disrupting the *nhj-1* sequence induces IR-sensitivity in an IR-resistant background. Double mutants of *nhj-1* and the cNHEJ factors *lig-4* or *cku-80* do not exhibit additive IR-sensitivity, arguing that *nhj-1* is a member of the cNHEJ pathway. Furthermore, like the loss of *lig-4,* the loss of *nhj-1* in the *com-1* genetic background, in which meiotic DSBs are repaired by cNHEJ instead of homologous recombination, increased the number of DAPI-staining bodies in diakinesis, consistent with increased chromosome fragmentation in the absence of cNHEJ repair. Finally, we show that NHJ-1 localizes to many somatic nuclei in the L1 larva, but not the primordial germline, which is in accord with a role in the predominantly somatically active cNHEJ. Although *nhj-1* shares no sequence homology with other known eukaryotic cNHEJ factors and is taxonomically restricted to the Rhadbitid family, its discovery underscores the evolutionary plasticity of even highly conserved pathways, and may represent a springboard for further characterization of cNHEJ in *C. elegans*.

## Introduction

Canonical non-homologous end joining (cNHEJ) is one of the two major DNA double-strand break (DSB) repair modalities, standing in contrast to homologous recombination repair (HRR) [1]. Although commonly described as error prone [2], cNHEJ is a very rapid and efficient mode of DSB repair, and is the preferred repair pathway in contexts where the fidelity of repair is less important than the imperative of restoring chromosome integrity, or whenever an appropriate repair template is unavailable, such as in somatic cells prior to S-phase [3, 4]. A number of “alternative” end joining pathways (Alt-EJ), including microhomology-mediated end joining and polymerase theta-mediated end joining [5, 6], have been identified, but these pathways appear to primarily, although not exclusively, act as “backup” DSB repair pathways in cNHEJ- and HR-deficient conditions [7].

The mechanism of cNHEJ involves three distinct steps: 1) DSB detection and tethering; 2) DNA end processing; and 3) terminal ligation (reviewed in [8–10]). The first step is effected by nearly universally conserved Ku ring, a heterodimeric complex composed of Ku70 and Ku80, efficiently detects and binds free DNA ends in a sequence-independent manner, stabilizing and protecting them from extensive resection [9]. It then acts as a “toolbelt” for cNHEJ [11], recruiting a complex of proteins which in mammals includes kinases/phosphatases (DNA-PKcs, PNKP), nucleases (Artemis, aprataxin, APLF), polymerases (Pol X family members), and helicases (WRN) [10]. These enzymes act in the second step of cNHEJ to process the free DNA ends into a ligation-compatible form [9]. Finally, the conserved DNA ligase LIG4, in complex with the structural protein XRCC4, which oligomerizes to form long filaments with its paralogs XLF and the recently discovered PAXX [1], performs the third, terminal ligation step, restoring chromosomal integrity [9].

Canonical non-homologous end joining is a widely conserved DSB repair pathway, and is present in all three domains of life [12]. Vertebrates possess the best studied cNHEJ system and the highest number of participating proteins [10], with the cNHEJ systems in other organisms studied so far conserving a subset of the vertebrate cNHEJ factors. The Ku ring and DNA Ligase IV are universally conserved [12–16], but DNA-PKcs is absent in flies, yeast, plants, and nearly all basal multicellular eukaryotic lineages [10, 13–15]. An Artemis ortholog exists in budding yeast but doesn’t participate in cNHEJ [14], but flies appear to have lost this gene [15]. Three Artemis orthologs exist in the *Arabidopsis* genome, but it is not clear whether they play a role in cNHEJ [13]. Orthologs of the mammalian cNHEJ scaffold proteins XRCC4 and XLF exist in *S. cerevisiae* and participate in cNHEJ [14]. *Arabidopsis* possesses only XRCC4 [13], and although *D. melanogaster* retains one XRCC4 ortholog and two XLF orthologs, it is not known whether they play a role in cNHEJ in the fruit fly [15]. By contrast to other eukaryotes, the cNHEJ system in the nematode *C. elegans* consists of only the Ku70/Ku80 orthologs CKU-70 and CKU-80, and the LIG4 ortholog LIG-4 [17]. While a homolog of WRN helicase exists, it plays no role in cNHEJ in the worm [17]. Thus, *C. elegans* has been hypothesized to either possess a “minimal” cNHEJ system, in which the Ku ring and LIG-4 are sufficient to repair the breaks, or to contain other factors which would be functionally analogous if not homologous to the cNHEJ factors in other organisms [16].

Here, we report the discovery of a fourth cNHEJ factor in *C. elegans*, which we named NHJ-1 (<u>n</u>on-<u>h</u>omologous end joining 1). It is encoded by *H19N07.3*, a gene which was found to confer resistance to bleomycin, a common chemotherapeutic drug that is radiomimetic (scb-1; <u>s</u>ensitive to <u>c</u>hemotherapeutic <u>b</u>leomycin 1) [18, 19]. Lacking homology to proteins outside the Rhabditid family or any conserved domains, we show that NHJ-1 is nevertheless essential for cNHEJ both in the L1 larva and in the adult germline, which is in accord with the reported bleomycin sensitivity. Thus, *C. elegans* appears to have reorganized an ancient and conserved DSB repair pathway by the addition of at least one taxonomically restricted protein and may consequently possess a larger cNHEJ toolkit than previously thought.

## Results

### An N2 strain variant exhibits unexpected ionizing radiation sensitivity

During the course of an RNAi screen to identify novel factors that modulate the response of the adult germ line in *C. elegans* following exposure to ionizing radiation (IR) during the L1 stage, we observed a striking difference in IR sensitivity between our wild-type N2 strain and the N2 from the neighboring laboratory of Dr. Richard Roy. Irradiated at the L1 stage with 75 Gy, Zetka lab N2 animals produced significantly fewer progeny than Roy lab N2 (**Figure S1**). N2 is the most commonly used wild-type strain used by the *C. elegans* research community and is assumed to be isogenic [20], which made the difference in the IR response surprising. Although the N2 strain from the <u>C</u>aenorhabditis <u>G</u>enetics <u>C</u>enter (CGC) showed the same post-IR brood size reduction as Zetka lab N2 (**Figure S1**), several N2 strains from other laboratories, as well as 18 non-N2 wild-type isolates of *C. elegans* and two isolates of *C. briggsae* displayed a resistant IR response (**Table S1**), suggesting that IR sensitivity arose in the N2 lineage at some point during its cultivation as a laboratory strain. We were interested in uncovering the origin of this IR sensitivity. To maintain a high degree of isogeneity, we derived a sensitive strain (henceforth, N2 [S]) from a single individual from the CGC N2, and a resistant strain (henceforth, N2 [R)] from a single animal isolated from the N2 strain from Eric Andersen’s laboratory (Northwestern University). All subsequent characterization of the phenomenology was performed in these two strains.

We first asked whether the IR response is dose dependent, or represents a binary response. Both N2 [S] and N2 [R] showed a dose-dependent response, with the brood size of N2 [R] decreasing from a median of 291.5 progeny in unirradiated animals to 256 progeny when treated with 25 Gy, 41 progeny when irradiated with 50 Gy, and 1 progeny when exposed to 75 Gy (Figure 1A). The unirradiated brood size of 284.5 progeny in N2 was not significantly reduced when exposed to 25 Gy (median 289 progeny) or 50 Gy (255 progeny), but was significantly reduced when irradiated at 75 Gy (median 191 progeny, p<0.001 vs unirradiated controls) (Figure 1A). At each tested IR dose, however, the brood size of N2 [S] animals was significantly lower than that of N2 [R] animals (p<0.05 at 25 Gy; p<0.001 at 50 Gy and 75 Gy) (Figure 1A).

**Figure 1.**
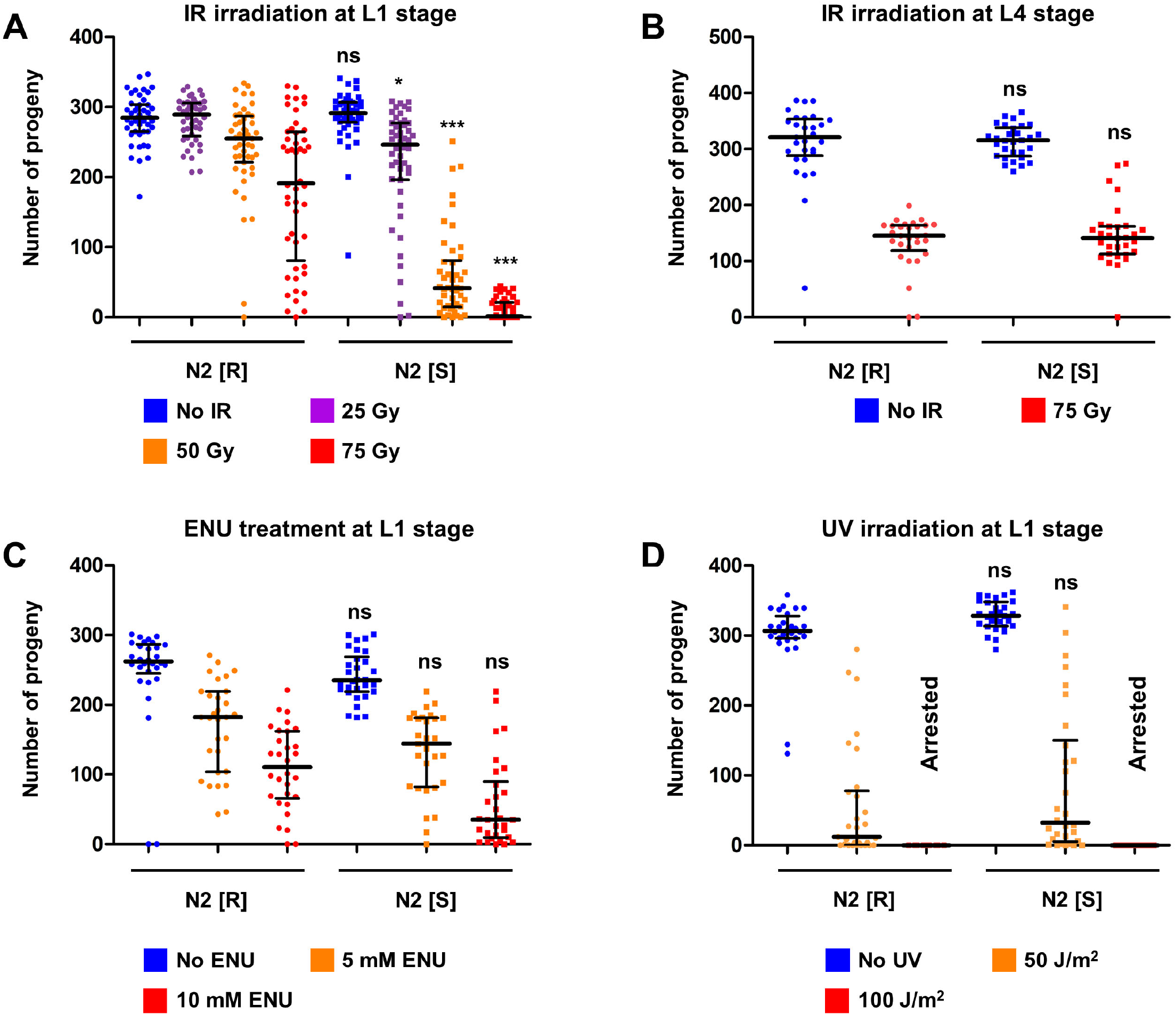
N2 [R] and N2 [S] show a dose-dependent, stage- and stressor-specific difference in brood size following treatment with ionizing radiation. **(A)** Multi-dose brood size quantification of N2 [R] and N2 [S] post-L1 IR treatment. The IR-sensitive N2 [S] line shows a significantly reduced brood size compared to unirradiated animals at 25 Gy (p<0.01), 50 Gy (p<0.001), and 75 Gy (p<0.001). Irradiated N2 [R] animals show a significantly reduced brood size compared to unirradiated controls only at 75 Gy (p<0.001). At every tested IR dose, N2 [S] is significantly more severely affected than N2 [R]. All statistical comparisons shown in the figure are to N2 [R] at the equivalent IR dose (Kruskal-Wallis test and Dunn’s post-hoc tests). Error bars represent the median and interquartile range. Sample size (n) is 46 for unirradiated N2 [R], 45 for N2 [R] 25 Gy, 47 for N2 [R] 50 Gy, 48 for N2 [R] 75 Gy, 44 for unirradiated N2 [S], 48 for N2 [S] 25 Gy, 46 for N2 [S] 50 Gy, and 49 for N2 [S] 75 Gy. **(B)** Brood size quantification of N2 [R] and N2 [S] post-L4 IR treatment. Both N2 [S] and N2 [R] animals show a reduced brood size in adulthood (p<0.001 vs unirradiated controls), but there is no significant difference between the two backgrounds, suggesting that the IR-sensitivity of N2 [S] is specific to the L1 stage. All statistical comparisons shown in the figure are to N2 [R] at the equivalent IR dose (Kruskal-Wallis test and Dunn’s post-hoc tests). Error bars represent the median and interquartile range. Sample size (n) is 29 for both N2 [R] groups, and 30 for both N2 [S] groups. **(C)** Brood size quantification of N2 [R] and N2 [S] post-L1 ENU treatment. Both N2 [S] and N2 [R] animals show a dose-dependent reduction in brood size (p<0.01 for 5mM ENU and p<0.001 for 10mM ENU for N2 [R] versus untreated control; p<0.001 for both 5mM and 10mM ENU for N2 [S] versus untreated control), but there is no significant difference between the two backgrounds at the doses tested. All statistical comparisons shown in the figure are to N2 [R] at the equivalent ENU dose (Kruskal-Wallis test and Dunn’s post-hoc tests). Error bars represent the median and interquartile range. Sample size (n) is 29 for N2 [S] 5mM ENU, and 30 for all other groups. **(D)** Brood size quantification of N2 [R] and N2 [S] post-L1 UV treatment. Both N2 [S] and N2 [R] animals show a reduction in adult brood size (p<0.001 versus unirradiated controls for both genotypes), but there is no significant difference between the two backgrounds. At the higher of the two tested doses, 100 J/m^2^, animals of both backgrounds exhibit terminal larval arrest and do not produce progeny. All statistical comparisons shown in the figure are to N2 [R] at the equivalent ENU dose (Kruskal-Wallis test and Dunn’s post-hoc tests). Error bars represent the median and interquartile range. Sample size (n) is 28 for unirradiated N2 [R], 29 for unirradiated N2 [S], and 30 for all other groups. ns = not significant (p>0.05); * = p<0.05; ** = p<0.01; *** = p<0.001

We then asked whether the difference in the IR response is general or developmentally restricted. To test this, we irradiated the animals at the L4 stage, when the somatic development is largely complete and the germline consists of hundreds of cells [21]. While irradiation with 75 Gy at the L4 stage did reduce the brood size in N2 [R] from a median of 321 progeny to a median of 145 progeny (p<0.001) and in N2 [S] from a mean of 315.5 progeny to a median of 141.5 progeny (p<0.001), there was no statistically significant difference between the two strains (Figure 1B). The reduction in brood size following IR at the L4 stage can be attributed to high embryonic lethality (>50% in both genotypes) (**Table S2**). Interestingly, embryonic lethality in N2 [S] animals irradiated with 75 Gy at L1 was not different than that of unirradiated controls, while irradiated N2 [R] animals showed a significantly increased embryonic lethality (9.45%) compared to unirradiated controls (0.63%, p<0.001) (**Table S2**).

We also wanted to test whether the differential brood size response of N2 [R] and N2 [S] is specific to IR or reflects a general difference in response to genotoxic stress. Treatment with ethyl-nitrosourea (ENU) which primarily causes base alkylation [22], resulted in a brood size reduction in both N2 [R] (untreated control median of 262 progeny, 5mM ENU median of 182 [p<0.01 vs untreated] and 10mM ENU median of 110.5 progeny [p<0.001 vs untreated] and N2 [S] (control median of 235 progeny, 5mM ENU median of 144 [p<0.001 vs untreated], and a 10mM ENU median of 35 progeny [p<0.001 vs untreated]) (Figure 1C). Despite a trend for lower brood size in N2 [S] at 10 mM ENU, there was no significantly different difference between the two genotypes at the doses tested. Irradiation with ultraviolet (UV) light, which predominantly causes pyrimidine dimers [23], likewise resulted in the same extent of brood size reduction at 50 J/m^2^ in both N2 [R] (unirradiated median of 306.5 progeny vs an irradiated median of 12 progeny [p<0.001]) and N2 [S] (control median of 328 progeny and an irradiated median of 32.5 progeny [p<0.001]), but there was no significant difference between the two strains. At 100 J/m^2^, UV treatment resulted in terminal larval arrest in both genotypes (Figure 1D). These data suggest that the sensitivity of N2 [S] is restricted to IR.

### N2 [S] displays somatic post-IR phenotypes

In addition to the difference in post-IR brood size, we noticed also noticed that N2 [S] animals displayed several post-IR somatic phenotypes that were observed with much lower frequencies in N2 [R] (Figure 2A-D). Prominently, irradiated N2 [S] animals exhibited a marked slow growth (Gro) phenotype. At 3 days after radiation treatment, when all control animals have developed into adults, as did 94% of irradiated N2 [R] animals, 61% of irradiated of the irradiated N2 [S] population was still in a larval stage, mostly (54%) L4 (p<0.001 vs irradiated N2 [R]) (Figure 2A). Furthermore, 27% of N2 [S] animals exhibited vulval phenotypes, including protruding vulva (Pvl) and ruptured through vulva (Rup), compared to just under 2% of irradiated N2 [R] animals (p<0.001) (Figure 2A). Four days after treatment, although the majority (88%) of irradiated N2 [S] animals developed into adults, the incidence of vulval phenotypes remained high (57% Pvl and 19% Rup) compared to that in irradiated N2 [R] (just under 5%; p<0.001) (Figure 2B), suggesting that the N2 [S] larvae were developing into morphologically abnormal adults. Consistent with the vulval phenotypes, a significantly higher proportion of irradiated N2 [S] animals (25-31%) exhibited an egg laying defective (Egl), compared to irradiated N2 [R] animals (2-3%; p<0.001) (Figure 2D). These phenotypes were reminiscent of those reported for irradiated cNHEJ mutants [17], suggesting the possibility that N2 [S] was IR sensitive because of a loss of function in cNHEJ.

**Figure 2.**
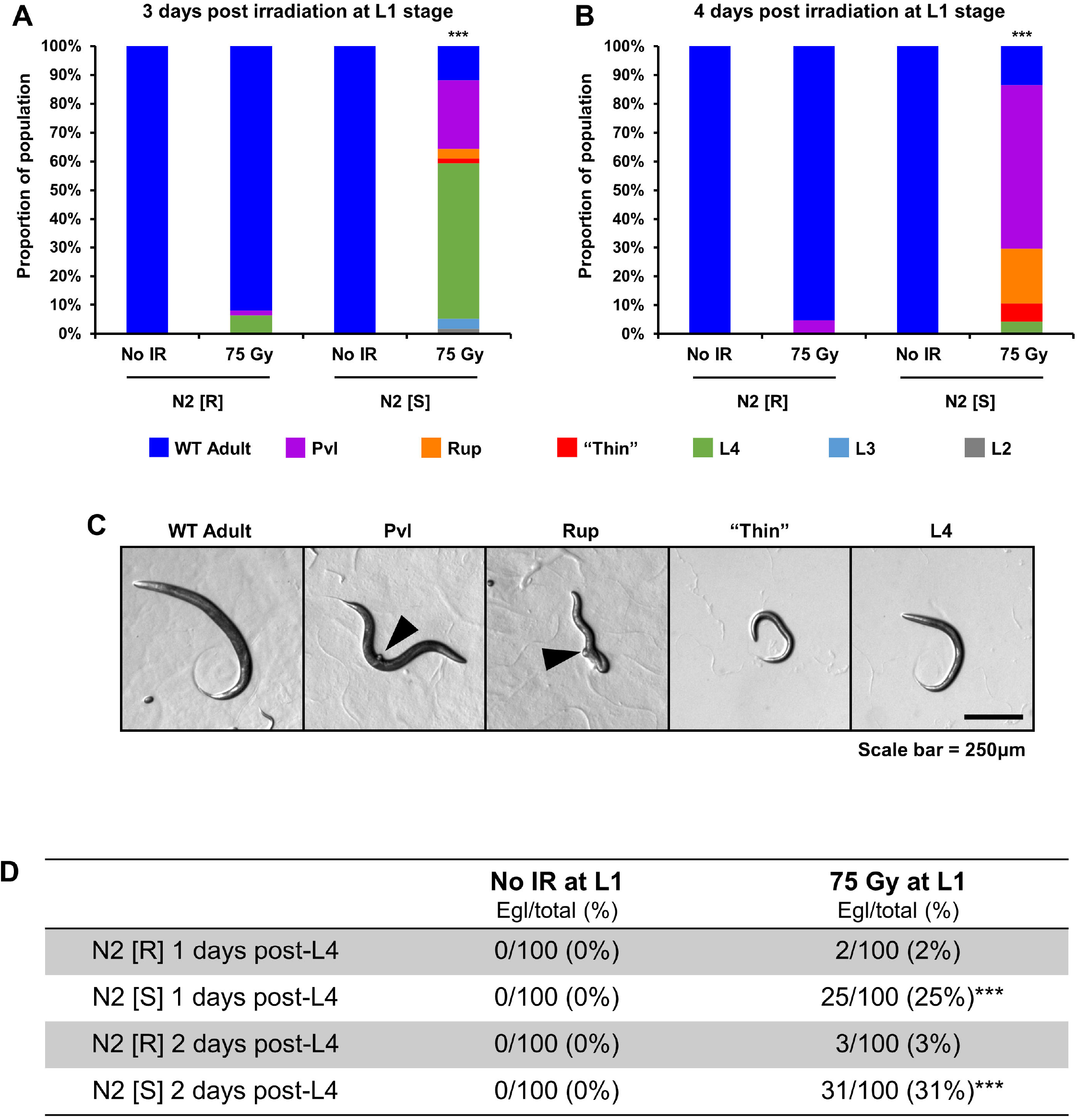
N2 [S] displays several distinct somatic phenotypes post-IR. **(A)** Quantification of growth delay and vulval phenotypes three days after IR treatment at the L1 stage. When irradiated at 75 Gy of IR, N2 [S] animals show a developmental delay, as a large proportion is still in the L4 stage when all unirradiated controls and almost all irradiated N2 [R] animals have developed into adults. A small proportion of irradiated N2 [S] animals develops into thin, whitish larvae approximately the size of L3 larvae (Thin). The majority of irradiated N2 [S] animals that does develop into adults by this stage exhibits vulval phenotypes, most prominently protruding vulva (Pvl) and ruptured through vulva (Rup). The statistical comparison shown in the figure is to irradiated N2 [R] (Chi-squared test). Sample size (n) is 148 for unirradiated N2 [R], 111 for irradiated N2 [R], 141 for unirradiated N2 [S], and 118 for irradiated N2 [S]. **(B)** Quantification of the same phenotypes as in **(A)**, four days after treatment at L1 stage. Four days after irradiation with 75 Gy of IR, almost all N2 [S] animals develop into adults, but exhibit a high incidence of Pvl and Rup phenotypes, as well as occasional thin, whitish larvae. The statistical comparison shown in the figure is to irradiated N2 [R] (Chi-squared test). Sample size (n) is 109 for unirradiated N2 [R], 148 for irradiated N2 [R], 134 for unirradiated N2 [S], and 142 for irradiated N2 [S]. **(C)** Representative images of phenotypes quantified in (A) and (B). Black arrows point to the protruding vulva in the “Pvl” panel and the burst vulva and partial extrusion of internal organs in the “Rup” panel. **(D)** Quantification of the Egl phenotype in irradiated and control animals. While a small proportion of N2 [R] animals develops the Egl phenotype, it is significantly more common in N2 [S] animals. The statistical comparisons shown in the figure are to irradiated N2 [R] at the equivalent time points (Chi-squared test). N2 [R] = resistant N2 strain, derived from Andersen lab N2 ns = not significant (p>0.01); * = p<0.05, *** = p<0.001

### N2 [S] is recessive to N2 [R]

A loss-of-function in a cNHEJ or another DNA repair factor would generally be expected to be genetically recessive. To test whether IR resistance is dominant or recessive to IR sensitivity, we compared the L1 IR response in F1 animals obtained from N2 [S] hermaphrodites mated to N2 [S] males, the N2 [S/S] homozygotes, and N2 [S] hermaphrodites mated to N2 [R] males, the N2 [R/S] heterozygotes. Irradiated with 50 Gy of IR at the L1 stage, the N2 [R/S] heterozygotes had a significantly higher brood size (median of 276 progeny) compared to N2 [S/S] homozygotes (median of 24 progeny; p<0.001) (Figure 3A). N2 [R/S] heterozygotes also exhibited a much lower frequency of somatic phenotypes four days after IR treatment, with a vulval phenotype incidence of just over 5%, compared to 62% in N2 [S/S] homozygotes (p<0.001) (Figure 3B), demonstrating that IR resistance is dominant over IR sensitivity, and suggesting the presence of a loss of function mutation(s) in the N2 [S] genetic background.

**Figure 3.**
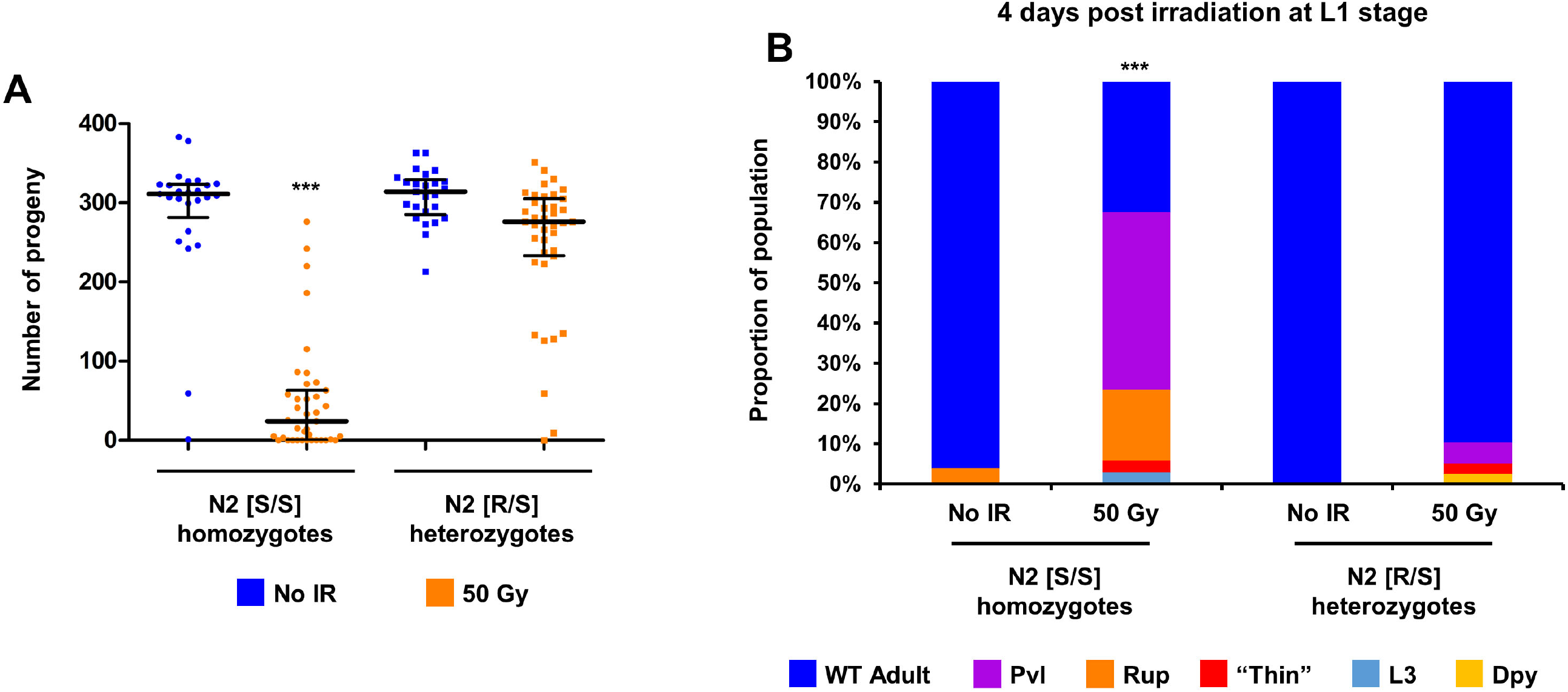
IR-resistance is dominant to IR-sensitivity. **(A)** Total brood size quantification in N2 [S/S] homozygotes and N2 [R/S] heterozygotes after 50 Gy of IR at the L1 stage. N2 [S/S] show a significantly lower post-IR brood size than N2 [R/S] heterozygotes, demonstrating that IR-sensitivity is a recessive trait. The statistical comparison shown in the figure is to irradiated N2 [R/S] heterozygotes (Kruskal-Wallis test, followed by Dunn’s post-hoc test). Error bars represent the median and interquartile range. Sample size (n) is 25 for both unirradiated groups and 39 for both irradiated groups. **(B)** Quantification of somatic phenotypes four days after IR treatment at the L1 stage. Irradiated N2 [S/S] homozygotes show a significantly higher incidence of Pvl and Rup phenotypes than N2 [R/S] heterozygotes, corroborating the conclusion that IR-sensitivity is recessive to IR-resistance. The statistical comparison shown in the figure is to irradiated N2 [R] (Chi-squared test). Sample size (n) is 25 for both unirradiated groups, 34 for irradiated N2 [S/S] homozygotes, and 39 for irradiated N2 [R/S] heterozygotes. *** = p<0.001

### Loss of *H19N07.3/nhj-1* function results in IR sensitivity

To identify the mutation causative of IR sensitivity in N2 [S], we sequenced the N2 [S] and N2 [R] genomes to high (>100X) coverage. Using coding variants unique to the N2 genome (**Table S3**) as molecular markers, we mapped (**Figure S2**) the causative locus to a region on chromosome V which contained an indel in the gene *H19N07.3*. Because of the resemblance of the IR-sensitivity of N2 [S] to that of cNHEJ mutants [17], we have named the gene *nhj-1* (<u>n</u>on-<u>h</u>omologous end joining 1). The N2 [S] mutation in *nhj-1* is an indel in exon 3 (Figure 4A), consisting of a 5-nucleotide deletion, an insertion of 107 nucleotides of unknown origin that are predicted to form a strong hairpin (Figure 4B), and an 8-nucleotide duplication. We have designated this allele *vv148*, and used CRISPR mutagenesis to create a 7-nucleotide deletion in *nhj-1* in the N2 [R] background (Figure 4A), designated *vv144.* Introducing the *nhj-1(vv144)* allele into the [R] background resulted in a significant reduction in fertility post-IR, with the median brood size of 4 progeny, compared to 179.5 progeny in N2 [R] (p<0.001) and 0 progeny in N2 [S] (p>0.05) (Figure 5A). All irradiated *nhj-1(vv144)* animals also exhibited either developmental delay or vulval phenotypes, in contrast to only 18% of N2 [R] animals which showed slowed growth (p<0.001), but not different than N2 [S] animals (p>0.05) (Figure 5B). This result strongly suggested that *nhj-1* is causative of the IR sensitivity in N2 [S]. To investigate this possibility, we performed a complementation test between the N2 [S] and N2 [R] genomes and *nhj-1(vv144)* [R]. While the N2 [R] genome was able to fully complement *nhj-1(vv144)* [R] mutants for post-IR brood size (median of 149 progeny in the heterozygote, compared to 108 progeny in N2 [R] [p>0.05] and 2 progeny in N2 [S] [p<0.001]), the N2 [S] genome was not (median of 2 progeny in the heterozygote [p<0.001 vs N2 [R] and p>0.05 vs N2 [S]]) (Figure 5C). Similarly, the incidence of somatic phenotypes in *nhj-1(vv144)* [R]/*nhj-1(vv148)* [S] heterozygotes (81.2%) was not different than in homozygous *nhj-1(vv148)* [S] animals (79.6%; p>0.05), while that of *nhj-1(vv144)* [R]/*nhj-1(+)* [R] heterozygotes (6.0%) was not different than that of *nhj-1(+)* [R] homozygotes (18.2%; p>0.05) (Figure 5D).

**Figure 4.**
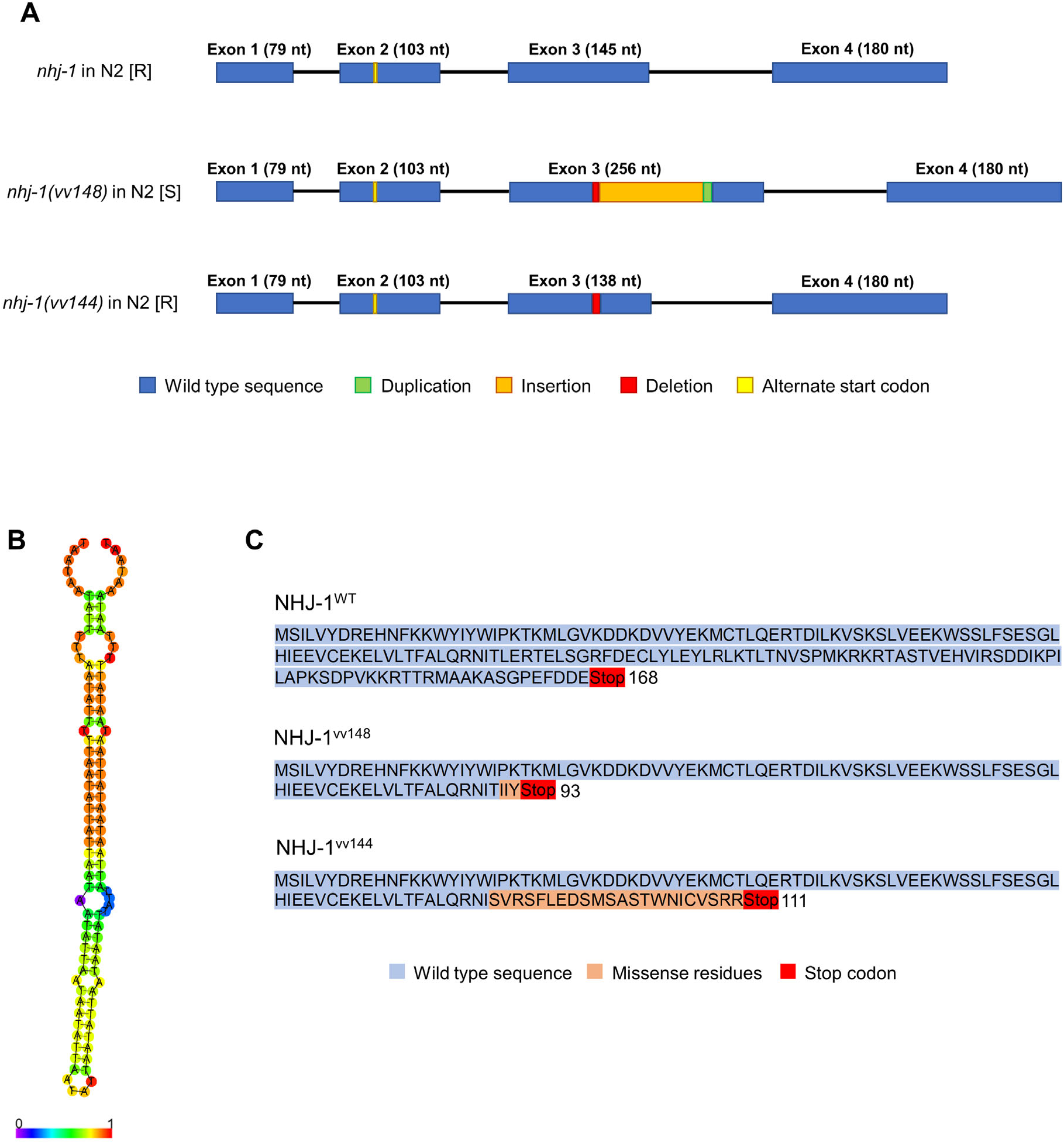
The structure of NHJ-1. **(A)** The structure of the coding region of *nhj-1*/*H19N07.3.* The uncharacterized gene *H19N07.3*, which I have named *nhj-1* (<u>n</u>on-<u>h</u>omologous end joining 1), is composed of four exonic regions and three introns. A shorter protein isoform can be translated from an alternate start codon in exon 2. In the N2 [S] background, exon 3 of *nhj-1* has been disrupted by a deletion of 5 nucleotides, and a 115 bp insertion composed of 107 nucleotides of unknown origin (see (**B)**) and 8 nucleotides duplicated from the exonic sequence. I have designated this mutation *nhj-1(vv148).* To test the role of *nhj-1* in IR-sensitivity, I used CRISPR mutagenesis to delete 7 nucleotides from Exon 3 and create the *nhj-1(vv144)* allele. **(B)** The predicted hairpin secondary structure of the 107 bp insertion in the *nhj-1(vv148)* allele using the RNAfold tool of the ViennaRNA Package. **(C)** The predicted protein sequences of NHJ-1. The wild-type long isoform of the NHJ-1 protein is 168 amino acids long, with no conserved domains. The shorter isoform is 130 residues in length. The *nhj-1(vv148)* indel results in truncated protein products of 93^long^/55^short^ amino acids in total length, with a frameshift producing 3 missense residues after residue 90^long^/52^short^. The *nhj-1(vv144)* deletion results in a frameshift after residue 89^long^/51^short^, which creates a downstream sequence of 22 missense residues before terminating in a stop codon, and produces final products 111^long^/73^short^ amino acids long.

**Figure 5.**
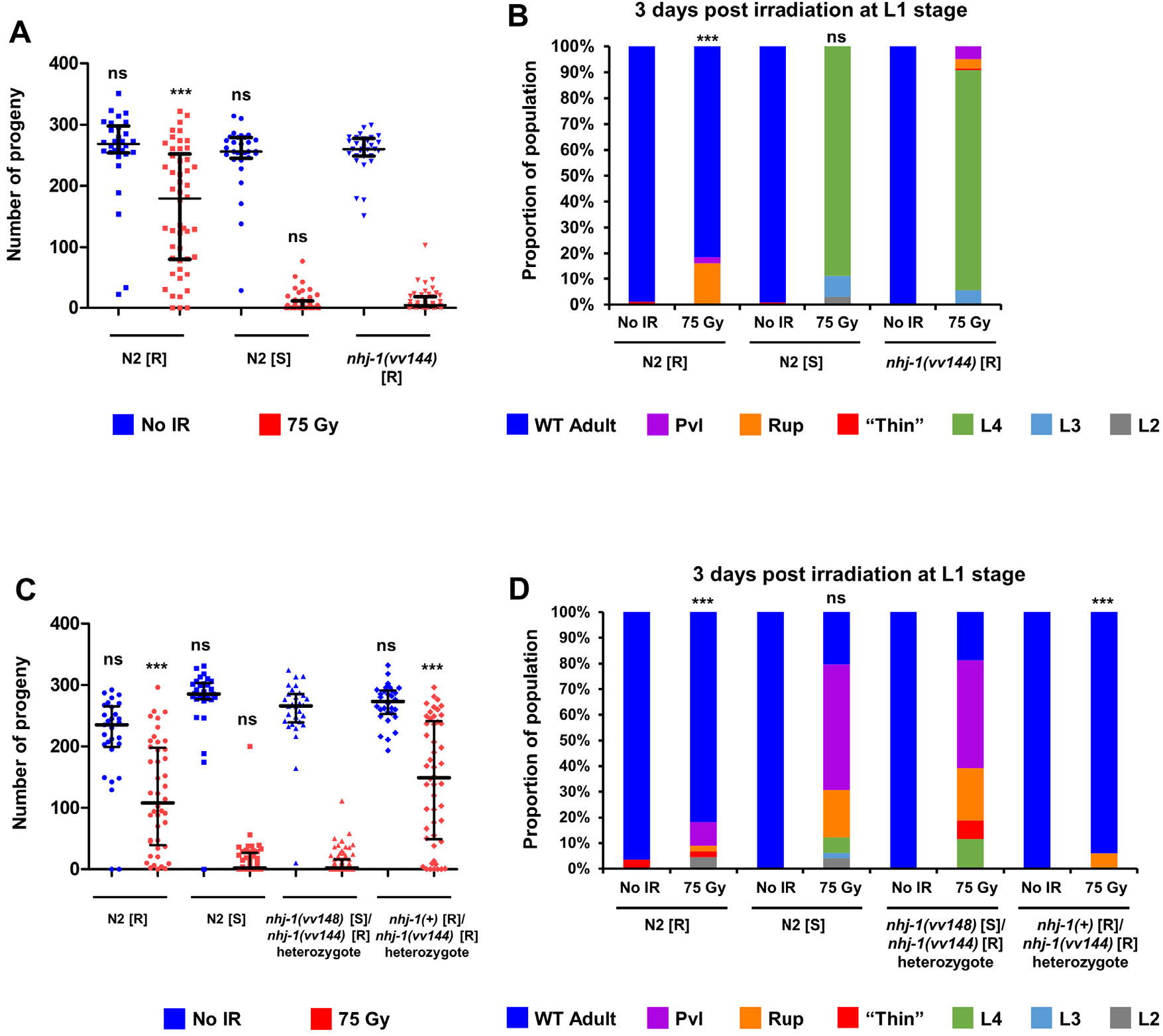
Loss of *nhj-1* in the N2 [R] background results in IR-sensitivity. **(A)** Total brood size quantification of N2 [R], N2 [S], and *nhj-1(vv144)*. While the *nhj-1(vv144)* deletion has no effect on untreated brood size (p>0.05 against both N2 [R] and N2 [S]), it significantly reduces the post-IR brood size of N2 [R] background compared (p<0.001 vs N2 [R] post-IR) to the same level as that of N2 [S] (p>0.05). All statistical comparisons shown in the figure are to *nhj-1(vv144)* from the corresponding treatment group (Kruskal-Wallis test, followed by Dunn’s post-hoc test). Error bars represent the median and interquartile range. Sample size (n) is 30 for all unirradiated groups, 50 for irradiated N2 [R] and irradiated *nhj-1(vv144)*, and 49 for irradiated N2 [S]. **(B)** Quantification of post-IR somatic phenotypes three days after IR treatment in the same groups as in **(A)**. Three days after IR treatment, *nhj-1(vv144)* mutants show a strong Gro phenotype, with almost all animals still in the L4 stage, like in N2 [S] (p>0.05), but significantly different than N2 [R] (p<0.001) in which all animals have molted into adults. All statistical comparisons shown in the figure are to irradiated *nhj-1(vv144)* (Chi-squared test, Bonferroni corrected for multiple comparisons to α = 0.01). Sample size (n) is 195 for unirradiated N2 [R], 161 for irradiated N2 [R], 252 for unirradiated N2 [S], 163 for irradiated N2 [S], 240 for unirradiated *nhj-1(vv144)*, and 163 for irradiated *nhj-1(vv144)*. **(C)** Total brood size quantification of N2 [R], N2 [S], and N2 [S]/*nhj-1(vv144)* and N2 [R]/*nhj-1(vv144)* heterozygotes. The post-IR brood size of N2 [S]/*nhj-1(vv144)* heterozygotes is not significantly different than that of post-IR N2 [S] animals (p>0.05), while both are significantly reduced compared to the post-IR brood size of either N2 [R] animals or N2 [R]/*nhj-1(vv144)* heterozygotes (p<0.001 for both comparisons), indicating that the IR-sensitivity of the N2 [S] line is caused by a loss of function in *nhj-*1. All statistical comparisons shown in the figure are to N2 [S]/*nhj-1(vv144)* heterozygotes from the corresponding treatment group (Kruskal-Wallis test, followed by Dunn’s post-hoc test). Error bars represent the median and interquartile range. Sample size (n) is 29 for unirradiated N2 [R], 44 for irradiated N2 [R], 30 for unirradiated N2 [S], 48 for irradiated N2 [S], 30 for unirradiated N2 [S]/*nhj-1* heterozygote, 68 for irradiated N2 [S]/*nhj-1(vv144)* heterozygote, 30 for unirradiated N2 [R]/*nhj-1* heterozygote, and 50 for irradiated N2 [R]/*nhj-1(vv144)* heterozygote. **(D)** Quantification of post-IR somatic phenotypes three days after IR treatment in the same groups as in **(C)**. The incidence of Gro and vulval phenotypes is not significantly different between N2 [S]/*nhj-1(vv144)* heterozygotes and N2 [S] animals after irradiation (p>0.05), while these phenotypes are significantly less common in post-IR N2 [R] animals and N2 [R]/*nhj-1(vv144)* heterozygotes (p>0.001 against both groups). All statistical comparisons shown in the figure are to N2 [S]/*nhj-1(vv144)* heterozygotes from the corresponding treatment group (Chi-squared test, Bonferroni corrected for multiple comparisons to α = 0.008). Sample size (n) is 28 for unirradiated N2 [R], 44 for irradiated N2 [R], 29 for unirradiated N2 [S], 49 for irradiated N2 [S], 30 for unirradiated N2 [S]/*nhj-1* heterozygote, 69 for irradiated N2 [S]/*nhj-1(vv144)* heterozygote, 30 for unirradiated N2 [R]/*nhj-1* heterozygote, and 50 for irradiated N2 [R]/*nhj-1(vv144).* ns = not significant (p>0.05 in **(A, C)**; p>0.01 in **(B)**; p>0.008 in **(D)**), *** = p<0.001

### Somatic and brood size IR phenotypes of *nhj-1* mutants are rescued by extrachromosomally expressed NHJ-1

We next wanted to test whether the loss of *nhj-1* function can be complemented molecularly by an extrachromosomal source of *nhj-1. nhj-1(vv144)* was crossed into a strain carrying *goeEx386*, a fosmid containing a GFP-tagged *nhj-1* sequence and the wild type *unc-119* sequence as a selection marker, which was created as part of the TransgeneOme project [24]. The post-IR brood size of *nhj-1(vv144); unc-119(ed3); goeEx386* animals (median of 131 progeny) was significantly higher than that of *nhj-1(vv144)* mutants alone (median of 1 progeny; p<0.001) (Figure 6A). Somatic phenotypes were also rescued in *nhj-1(vv144); unc-119(ed3); goeEx386*, with only 4.3% of animals showing Gro or vulval phenotypes, compared to 94.6% (p<0.001) in *nhj-1(vv144)* mutants (Figure 6B). These results suggest that both somatic and brood size phenotypes of *nhj-1(vv144)* can be rescued by an exogenous wild-type copy of the gene. To test this further, and to remove the potentially confounding effect of the rescue of *unc-119(ed3)*, we also compared the post-IR phenotypes of *nhj-1(vv148)* animals heterozygous for *unc-119* to those of the same genotype but also carrying the *goeEx386* fosmid. The post-IR brood size of animals with the extrachromosomal *nhj-1* was significantly higher (median of 109.5 progeny) than that of animals without the fosmid (median of 12.5 progeny, p<0.001) (Figure 6C). The incidence of somatic phenotypes is significantly lower in the animals with the fosmid (26.7%) than without (69.6%, p<0.001) (Figure 6D). These results collectively indicate that the loss of *nhj-1* activity leads to IR sensitivity.

**Figure 6.**
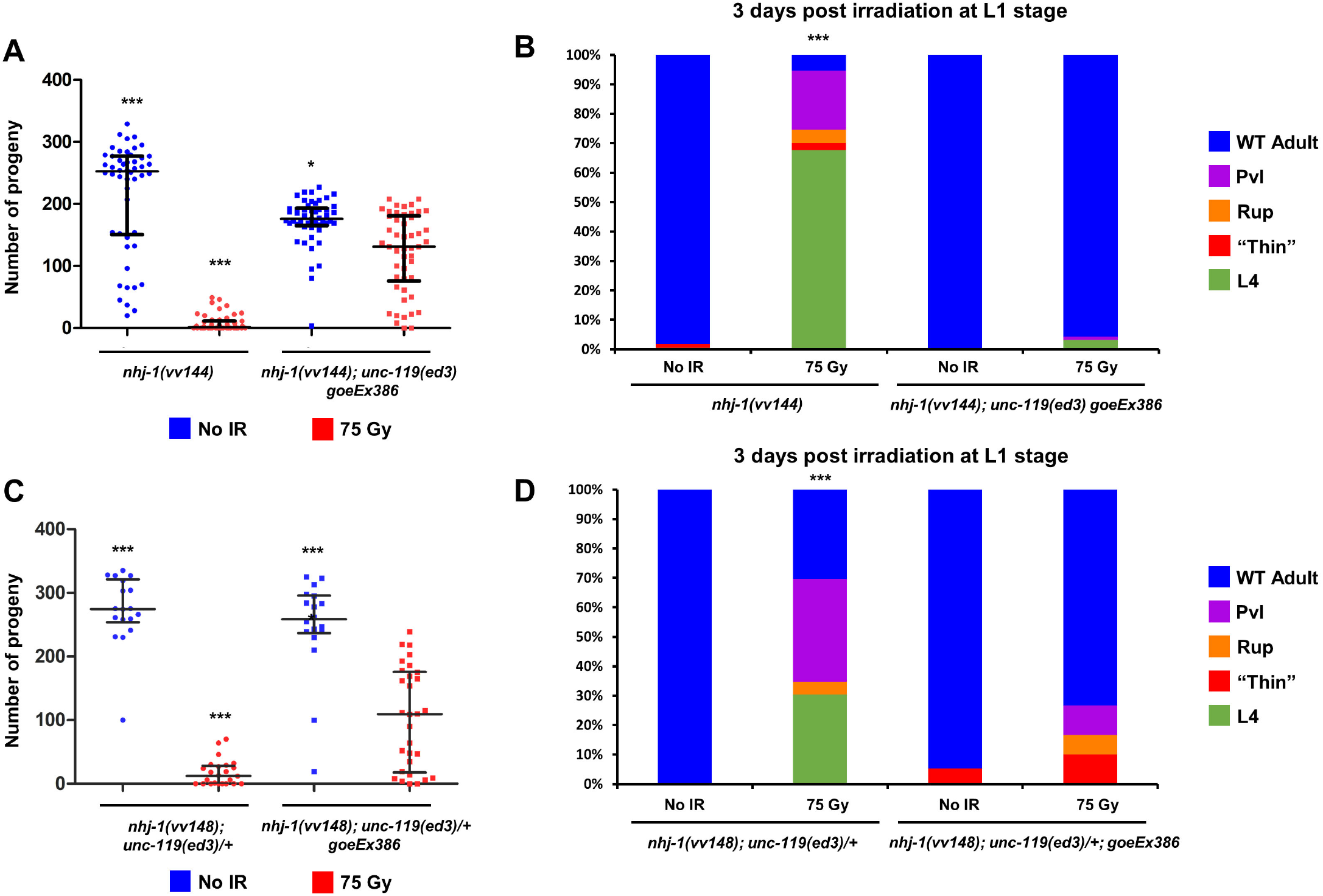
Extrachromosomal *nhj-1* rescues both brood size and somatic IR phenotypes of *nhj-1(vv144)* **(A)** Total brood size quantification of *nhj-1(vv144)* and *nhj-1(vv144); unc-119(ed3) goeEx386*. The post-IR brood size of *nhj-1(vv144); unc-119(ed3) goeEx386* animals, in which the *nhj-1(vv144)* allele and the *unc-119(ed3)* allele have been rescued by and extrachromosomal array carrying a GFP-tagged copy of wild type *nhj-1* and a wild type copy of *unc-119*, is significantly higher than that of *nhj-1(vv144)* animals (p<0.005), and is also reduced compared to unirradiated *nhj-1(vv144); unc-119(ed3) goeEx386* controls. Extrachromosomal *nhj-1* is thus able to rescue the post-IR brood size phenotype of *nhj-1(vv144).* All statistical comparisons shown in the figure are to irradiated *nhj-1(vv144); unc-119(ed3) goeEx386* animals (Kruskal-Wallis test, followed by Dunn’s post-hoc test). Error bars represent the median and interquartile range. Sample size (n) is 50 for all groups. **(B)** Quantification of post-IR somatic phenotypes three days after IR treatment in the same groups as in **(A)**. The Gro and vulval phenotypes are significantly less prevalent in irradiated *nhj-1(vv144); unc-119(ed3) goeEx386* animals compared to the irradiated *nhj-1(vv144)* group (<0.001), showing that exogenous *nhj-1* rescues the post-IR defects of *nhj-1(144)*. The statistical comparison shown in the figure is to irradiated *nhj-1(vv144); unc-119(ed3) goeEx386* animals (Chi-squared test, Bonferroni corrected for multiple comparisons to α = 0.008). Sample size (n) is 220 for unirradiated *nhj-1(vv144)* NR, 130 for irradiated *nhj-1(vv144)*, 70 for unirradiated *nhj-1(vv144); unc-119(ed3) goeEx386*, and 93 for irradiated *nhj-1(vv144); unc-119(ed3) goeEx386*. **(C)** Total brood size quantification of *nhj-1(vv148); unc-119(ed3)/+* and *nhj-1(vv148); unc-119(ed3)/+ goeEx386*. In *nhj-1(vv148); unc-119(ed3)/+ goeEx386* animals, the post-IR brood size is significantly rescued compared to *nhj-1(vv148); unc-119(ed3)/+* animals which do not carry the rescuing transgene (p<0.05), corroborating the conclusion that extrachromosomal *nhj-1* can rescue a lack of endogenous *nhj-1.* All statistical comparisons shown in the figure are to irradiated *nhj-1(vv148); unc-119(ed3)/+ goeEx386* animals (Kruskal-Wallis test, followed by Dunn’s post-hoc test). Error bars represent the median and interquartile range. Sample size (n) is 18 for both unirradiated groups, 24 for irradiated *nhj-1(vv148); unc-119(ed3)/+*, and 30 for irradiated *nhj-1(vv148); unc-119(ed3)/+ goeEx386*. **(D)** Quantification of post-IR somatic phenotypes three days after IR treatment in the same groups as in **(C)**. Vulval phenotypes and slow growth have a lower incidence in irradiated *nhj-1(vv148); unc-119(ed3)/+ goeEx386* animals compared to irradiated *nhj-1(vv148); unc-119(ed3)/+* animals (<0.001), in line with the brood size results. The statistical comparison shown in the figure is to irradiated *nhj-1(vv148); unc-119(ed3)/+; goeEx386* animals (Chi-squared test, Bonferroni corrected for multiple comparisons to α = 0.008). Sample size (n) is 18 for unirradiated *nhj-1(vv148); unc-119(ed3)/+*, 19 for unirradiated *nhj-1(vv148); unc-119(ed3)/+ goeEx386*, 23 for irradiated *nhj-1(vv148); unc-119(ed3)/+*, and 30 for irradiated *nhj-1(vv148); unc-119(ed3)/+ goeEx386*.n ns = not significant (p>0.05 in **(A)**; p>0.008 in **(B)**), * = p<0.05, *** = p<0.001

### NHJ-1 functions in the canonical non-homologous end joining pathway

To genetically test whether *nhj-1* has a role in the cNHEJ pathway, we used CRISPR mutagenesis to inactivate the terminal cNHEJ ligase, *lig-4*, in the N2 [R] and the N2 [S] background, hypothesizing that if *nhj-1* and *lig-4* function in different pathways, the double mutant would have a more severe phenotype than either single mutant. Because both N2 [S] (Figure 1A, Figure 2A-B) and the *lig-4(vv134)* null allele (**Figure S3**) in the resistant background exhibit a response so severe at 75 Gy that additivity would be difficult to discern, we halved the dose to 37.5 Gy. At this dose, the resistant and sensitive response are clearly distinguishable, but the sensitive response is not so severe as to not be augmentable. After treatment with 37.5 Gy of IR, *lig-4* mutants in the [S] background showed a median brood size of 170.5 progeny, which was not significantly different from either *lig-4* mutants in the [R] background (median brood size of 102 progeny; p>0.05) or N2 [S] animals (median brood size of 188 progeny; p>0.05), but significantly lower than that of N2 [R] animals (median brood size of 290 progeny; p<0.001) (Figure 7A). The incidence of somatic phenotypes in *lig-4* [S] animals (45.2%) was not significantly different than that of *lig-4* [R] animals (48.1%; p>0.05), but was higher than that of N2 [S] animals (25.5%; p<0.001) or N2 [R] animals (0.0%, p<0.001) (Figure 7B). Because no additivity in phenotype was observed between *lig-4* and the inactivation of *nhj-1* in the N2 [S] background, these results strongly suggest that *lig-4* and *nhj-1* fall in the same pathway, and that *nhj-1* is a member of the cNHEJ pathway. We also tested for additivity of IR response in double mutants carrying *nhj-1(vv144)* and a published allele of the Ku ring component, *cku-80(tm1203)*. The median post-IR brood size of *cku-80(tm1203); nhj-1(vv144)* double mutants was 40.5 progeny, which was not significantly different than either *cku-80(tm1203)* single mutants (median of 55 progeny; p>0.05) or *nhj-1(vv144)* single mutants (median of 70 progeny; p>0.05), but was significantly lower than that of N2 [R] animals (median of 187.5 progeny; p<0.001) (Figure 7C). Similarly, the somatic phenotype incidence of *cku-80(tm1203); nhj-1(vv144)* double mutants (40.1%) was not significantly different than *cku-80(tm1203)* single mutants (46.0%, p>0.05) or *nhj-1(vv144)* single mutants (45.5%; p>0.05), and higher than that of N2 [R] animals (6.2%, p<0.001) (Figure 7D). Collectively with the results of the *lig-4* double mutants, these observations support the interpretation that NHJ-1 acts in the cNHEJ pathway.

**Figure 7.**
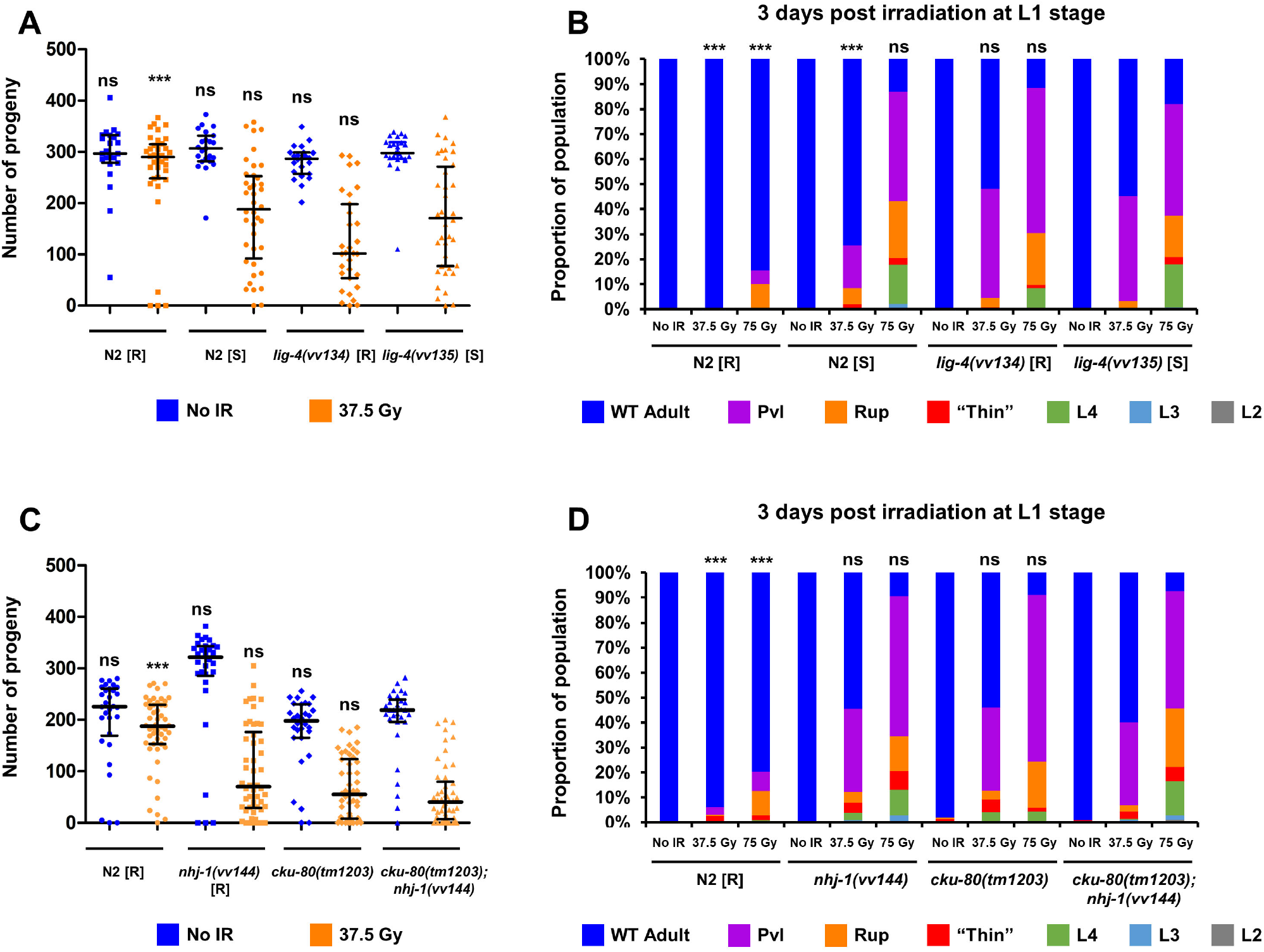
NHJ-1 acts in the cNHEJ pathway. **(A)** Total brood size quantification of N2 [R], N2 [S], *lig-4(vv134)* [R], and *lig-4(vv141)* [S] hermaphrodites. The post-IR brood size of *lig-4(vv141)* [S] animals, which harbour the same inactivating mutation as *lig-4(vv134)* [R] animals except in the sensitive genetic background, is not significantly different than either N2 [S] or *lig-4(vv134)* [R] animals (p>0.05). The lack of additive IR-sensitivity strongly suggests that N2 [S] is IR-sensitive because of a loss of cNHEJ activity. All statistical comparisons shown in the figure are to *lig-4* [S] animals from the corresponding treatment group (Kruskal-Wallis test, followed by Dunn’s post-hoc test). Error bars represent the median and interquartile range. Sample size (n) is 23 for unirradiated N2 [R], 36 for irradiated N2 [R], 23 for unirradiated N2 [S], 40 for irradiated N2 [S], 23 for unirradiated *lig-4* [R], 31 for irradiated *lig-4* [R], 24 for unirradiated *lig-4* [S], and 40 for irradiated *lig-4* [S]. **(B)** Quantification of post-IR somatic phenotypes three days after IR treatment in the same groups as in **(A)**. The incidence of somatic phenotypes in *lig-4* [S] is not significantly different from either *lig-4* [R] or N2 [S] following either 37.5 Gy or 75 Gy of IR (p>0.05 for all comparisons), showing that the *lig-4* mutation and the N2 [S] background are not additive with respect to IR-associated somatic phenotypes. All statistical comparisons shown in the figure are to *lig-4* [S] animals from the corresponding treatment group (Chi-squared test, Bonferroni corrected for multiple comparisons to α = 0.008). Sample size (n) is 146/201/181 for N2 [R] No IR/37.5 Gy/75 Gy, 188/106/146 for N2 [S] No IR/37.5 Gy/75 Gy, 208/131/167 for *lig-4* [R] No IR/37.5 Gy/ 75 Gy, and 177/134/125 for *lig-4* [S] No IR/37.5 Gy/ 75 Gy. **(C)** Total brood size quantification of N2 [R], *nhj-1(vv144)* [R], *cku-80(tm1203)*, and *cku-80(tm1203); nhj-1(vv144)*. Double mutants of *cku-80(tm1203)* and *nhj-1(vv144)* do not exhibit a significantly different post-IR brood size than either single mutant (p>0.05 for both), showing that NHJ-1 functions in the same pathway as CKU-80. All statistical comparisons shown in the figure are to *cku-80(tm1203); nhj-1(vv144)* animals from the corresponding treatment group (Kruskal-Wallis test, followed by Dunn’s post-hoc test). Error bars represent the median and interquartile range. Sample size (n) is 23 for unirradiated N2 [R], 36 for irradiated N2 [R], 23 for unirradiated N2 [S], 40 for irradiated N2 [S], 23 for unirradiated *lig-4* [R], 31 for irradiated *lig-4* [R], 24 for unirradiated *lig-4* [S], and 40 for irradiated *lig-4* [S]. **(D)** Quantification of post-IR somatic phenotypes three days after IR treatment in the same groups as in **(C)**. Vulval and slow growth phenotypes do not have a significantly different incidence in the double mutant and either single mutant (p>0.05 for all comparisons), supporting the conclusion of CKU-80 and NHJ-1 acting in the same pathway. All statistical comparisons shown in the figure are to *cku-80(tm1203); nhj-1(vv144)* animals from the corresponding treatment group (Chi-squared test, Bonferroni corrected for multiple comparisons to α = 0.008). Sample size (n) is 181/194/103 for N2 [R] No IR/37.5 Gy/75 Gy, 156/211/107 for *nhj-1(vv144)* [R] No IR/37.5 Gy/75 Gy, 156/163/136 for *cku-80(tm1203)* No IR/37.5 Gy/75 Gy, and 225/274/175 for *cku-80(tm1203); nhj-1(vv144)* No IR/37.5 Gy/75 Gy. ns = not significant (p>0.05 in **(A)**; p>0.008 in **(B)**), *** = p<0.001

### NHJ-1 acts downstream of Ku

Previously known *C. elegans* cNHEJ factors include the Ku ring components CKU-70 and CKU-80, which presumably act in DSB detection and DNA end protection/tethering, and LIG-4, which presumably performs the terminal ligation step of the pathway [1, 17]. NHJ-1 could be acting together with Ku in the first step, or at any of the downstream steps described in other organisms [8–10], including as a processing factor, a structural scaffold, or a co-factor in an enzymatic reaction. To test whether NHJ-1 acts in cNHEJ initiation, or at a downstream event, we made use of a *com-1* deficient background. In prophase I of *C. elegans* meiosis, COM-1 acts to prevent Ku binding to the free DNA ends of meiotic DSBs generated by the topoisomerase II-like enzyme SPO-11 [25]. This allows for inter-homolog HRR and crossover (CO) formation to take place, which is a critical both for the generation of allelic diversity and for proper segregation of homologous chromosomes in anaphase I [26]. Loss of COM-1 function thus results in cNHEJ-based repair of SPO-11 induced meiotic DSBs, leading to a loss of COs and nearly complete embryonic lethality [25]. The lethality of *com-1* mutants can be rescued by removal of either *cku-70* or *cku-80*, consistent with the idea that HRR can initiate on meiotic DSBs if the Ku ring has not loaded, but cannot be rescued by the loss of *lig-4*, consistent with the interpretation that Ku loading prevents HRR even if repair by cNHEJ cannot complete [25]. Consistent with previous reports, we found that RNAi-mediated knockdown of *cku-80* in the *com-1(t1626)* genetic background restores embryonic viability to 26.42%, compared to 0.31% in control *com-1(t1626)* animals (p<0.001) (Figure 8A). Also in agreement with published data, *com-1(t1626) lig-4(vv134)* double mutants have the same embryonic viability (0.53%) as *com-1(t1626)* single mutants (p>0.05) (Figure 8A). The embryonic viability of *com-1(t1626); nhj-1(vv144)* double mutants (0.00%) is also not significantly different than that of *com-1(t1626)* single mutants (p>0.05) (Figure 8A). The lack of rescue of embryonic viability thus suggests that *nhj-1* is dispensable for Ku loading to meiotic DSBs. The morphology of DAPI-staining bodies was diverse but similar in *com-1* single mutants and *com-1 lig-4* and *com-1; nhj-1* double mutants (Figure 8B). The number of DAPI-staining bodies in *com-1; nhj-1* double mutants (median of 5 bodies) was not significantly different than that of *com-1; lig-4* double mutants (median of 4 bodies, p>0.05), but was significantly higher than that of *com-1* single mutants (median of 3 bodies, p<0.001) (Figure 8C), consistent with the interpretation that a deficiency in cNHEJ activity in *lig-4* and *nhj-1* mutant backgrounds leads to greater chromosome fragmentation.

**Figure 8.**
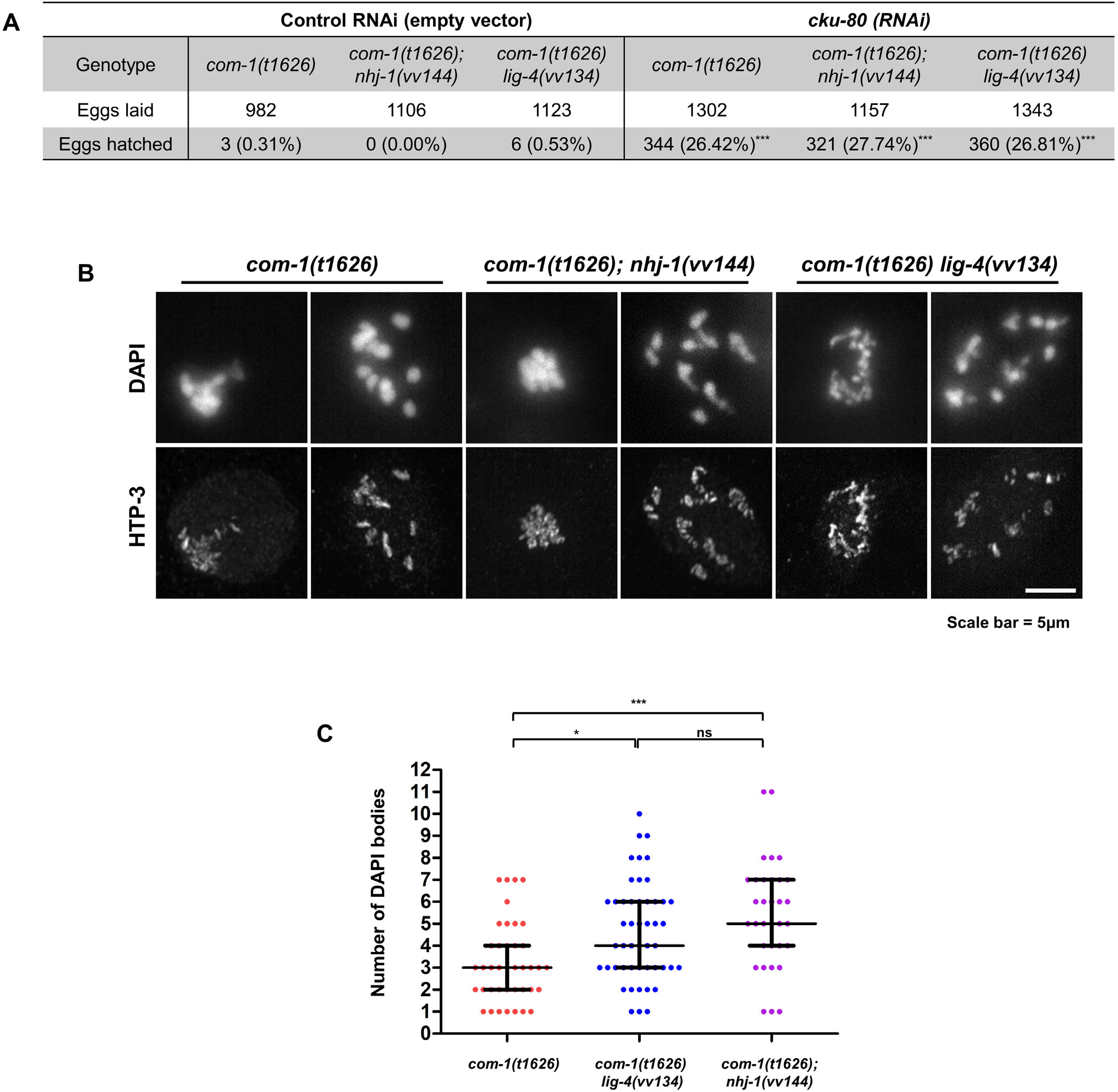
NHJ-1 acts downstream of the Ku ring in the adult germline. **(A)** Table showing the proportion of eggs hatching in *com-1, com-1; nhj-1,* and *com-1 lig-4* mutants treated with *cku-80(RNAi)* and controls. In control conditions, only a small fraction (<1%) of eggs laid in all three genotypes hatch. With RNAi against *cku-80*, the proportion of hatching eggs is significantly increased (p<0.001 versus RNAi control) in all three genotypes. All statistical comparisons shown in the figure are to the RNAi control group within the same genotype (Chi-squared test, Bonferroni correction for multiple comparisons to = 0.008). **(B)** Example micrographs showing the diverse DNA morphologies in diakinesis nuclei with low and high numbers of DAPI-staining entities in *com-1, com-1; nhj-1*, and *com-1 lig-4* mutants. **(C)** Quantification of DAPI-staining bodies in *com-1*, *com-1 lig-4*, and *com-1; nhj-1* mutants. The number of DAPI-staining bodies is significantly higher in *com-1 lig-4* (p<0.05) and *com-1; nhj-1* (p<0.001) double mutants is significantly higher than that of *com-1* single mutants, while the two double mutants are not significantly different from each other (p>0.05). Sample size (n) is 40 for *com-1*, 49 for *com-1 lig-4*, and 33 for *com-1; nhj-1* mutants.

### Endogenous NHJ-1 protein localizes to somatic cell nuclei in the L1 larva and meiocyte nuclei of prophase I in adult animals

We also wanted to investigate the localization pattern of the endogenous NHJ-1 protein, and tagged the C-terminus of *nhj-1* with the small epitope tag OLLAS [27]. Immunostaining with anti-OLLAS antibodies revealed that NHJ-1 localizes to many somatic cell nuclei in the L1 larva, but is conspicuously absent from the primordial germ cells (Figure 9A, 9B), consistent with a role in cNHEJ, which occurs in the nucleus and is primarily restricted to somatic cells [17]. Radiation treatment did not visibly alter this localization pattern (Figure 9A, 9B). Because in mammalian cNHEJ, the nuclear localization of XRCC4 depends on its binding partner LIG4 [28], we also wanted to test whether the localization of NHJ-1 depended on the Ku ring or LIG-4. However, the localization pattern of NHJ-1 is not affected in backgrounds deficient for either *cku-80* or *lig-4* (Figure 9A, 9B), demonstrating that NHJ-1 does not require these cNHEJ components for localization to the nucleus. In the adult germline, NHJ-1 first becomes reliably detectable in diplotene nuclei, and persists in diakinesis (Figure 10A), consistent with its role in cNHEJ, which in adult meiocytes is a backup DNA repair process that normally occurs only in the absence of functional COM-1. Like in the L1 larvae, the localization of NHJ-1 in the adult germline is unaffected by either the loss of *cku-80* (Figure 10B) or *lig-4* (Figure 10C).

**Figure 9.**
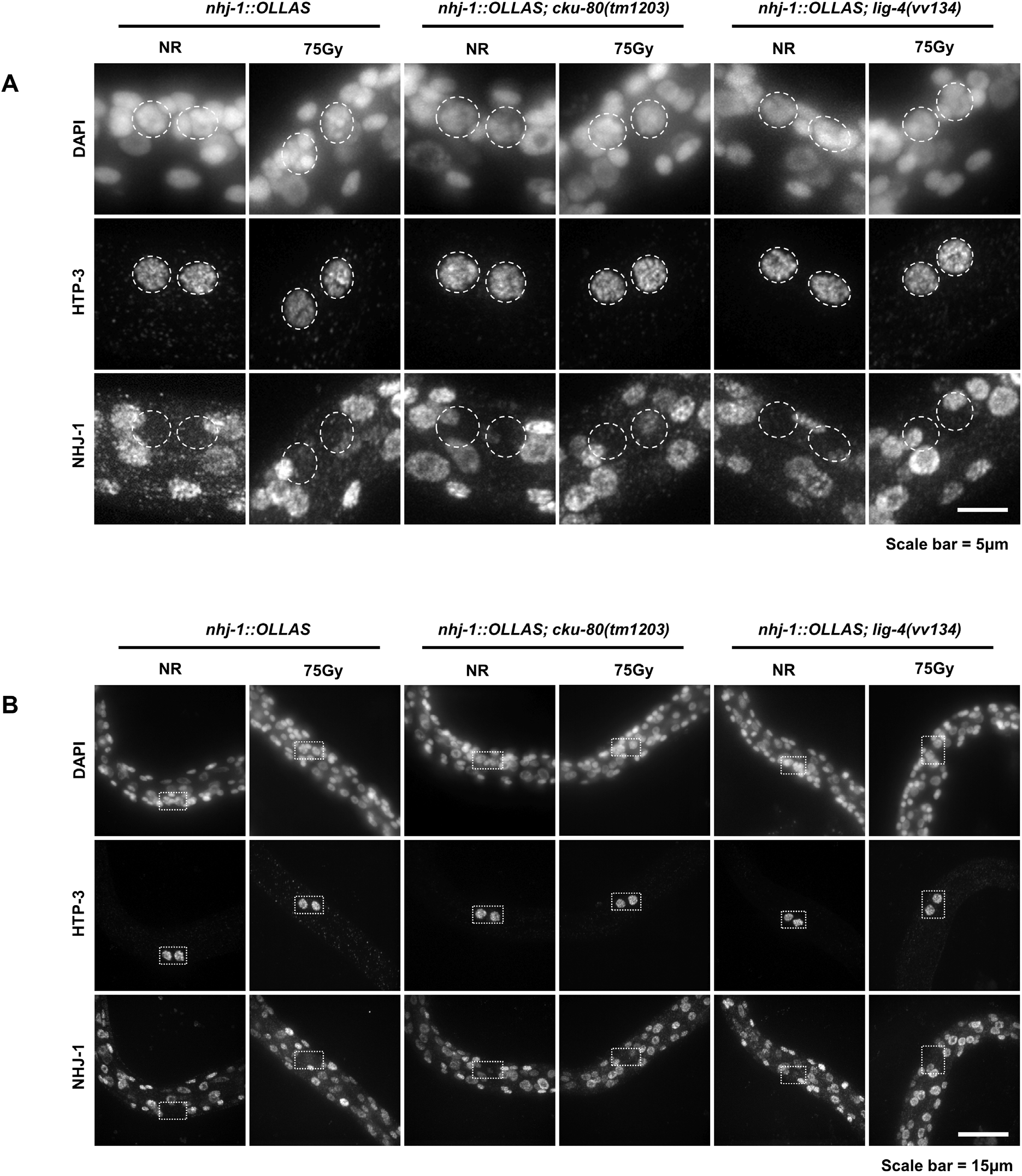
Endogenous NHJ-1 localization in the L1 larva. **(A)** Representative micrographs showing the subcellular localization of NHJ-1::OLLAS from the endogenous locus, together with DNA staining (DAPI) and the germline marker HTP-3, in the L1 larva. The loss of *cku-80* or *lig-4* does not detectably affect the localization of NHJ-1::OLLAS, and neither does the radiation treatment in either the control or *cku-80* or *lig-4* mutant backgrounds. Dotted lines delineate PGC nuclei. **(B)** Representative micrographs showing the subcellular localization of NHJ-1::OLLAS in L1 larvae in the same genotypes and conditions as in **(A)**, showing a wider field of view for comparison. Dotted lines box the PGCs.

**Figure 10.**
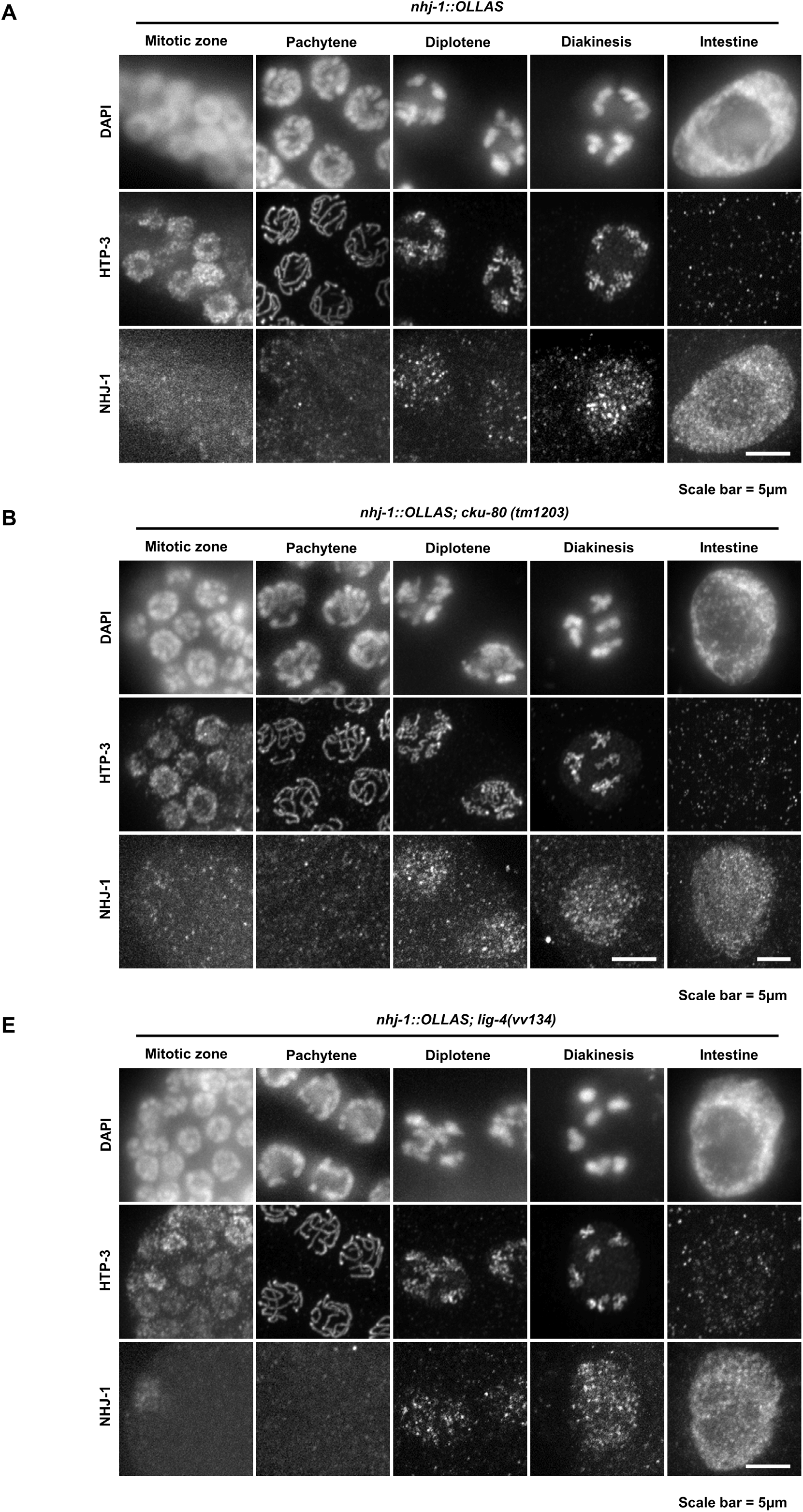
Endogenous NHJ-1 localization in the adult germline. **(A)** Representative micrographs of NHJ-1::OLLAS expression from the endogenous locus in adult germline cells. Punctate nuclear expression of NHJ-1::OLLAS becomes reliably visible in diplotene, but is not chromatin associated and remains detectable in diakinesis. Adult intestinal cell shown for comparison. **(B)** Representative micrographs of NHJ-1::OLLAS expression from the endogenous locus in adult germline cells in animals deficient for *cku-80.* The loss of CKU-80 does not perturb the localization of NHJ-1::OLLAS either in the germline or in the intestine. **(C)** Representative micrographs of NHJ-1::OLLAS expression from the endogenous locus in adult germline cells in animals deficient for *lig-4.* Like the loss of CKU-80, the loss of LIG-4 does not affect the pattern of NHJ-1::OLLAS expression either in the germ cells or intestinal cells.

### Endogenous LIG-4 localizes to intestinal cell nuclei in the L1 larva, and nuclei of adult prophase I meiocytes

Since nothing is known about the endogenous localization pattern of the previously described *C. elegans* cNHEJ factors, we also investigated the localization of LIG-4, the terminal effector of the pathway. Like NHJ-1, LIG-4 is not detectable in the primordial germ cells of the L1 larva (Figure 11A, 11B). Unlike NHJ-1, it is reliably detectable only in a longitudinal array of somatic cell nuclei (Figure 11A, 11B), which are inferred to be intestinal cell nuclei since they co-express a GFP reporter under the control of the intestinal cell promoter *elt-2* (Figure 11C). Like that of NHJ-1, the localization pattern of LIG-4 in the L1 is unaffected by the loss of other known cNHEJ factors, *nhj-1* and *cku-80*, or by exposure to ionizing radiation (Figure 11A, 11B). In the adult meiocytes, the localization of LIG-4 is similar to that of NHJ-1, except that it becomes reliably detectable already in pachytene, persisting into diakinesis (Figure 12A). Like in the L1, this pattern is not altered in absence of *cku-80* (Figure 12B) or *nhj-1* (Figure 12C). The distinct patterns of LIG-4 and NHJ-1 localization raise the question of now cNHEJ is coordinated in various tissue contexts, and whether individual components of the pathway may have pleiotropic roles.

**Figure 11.**
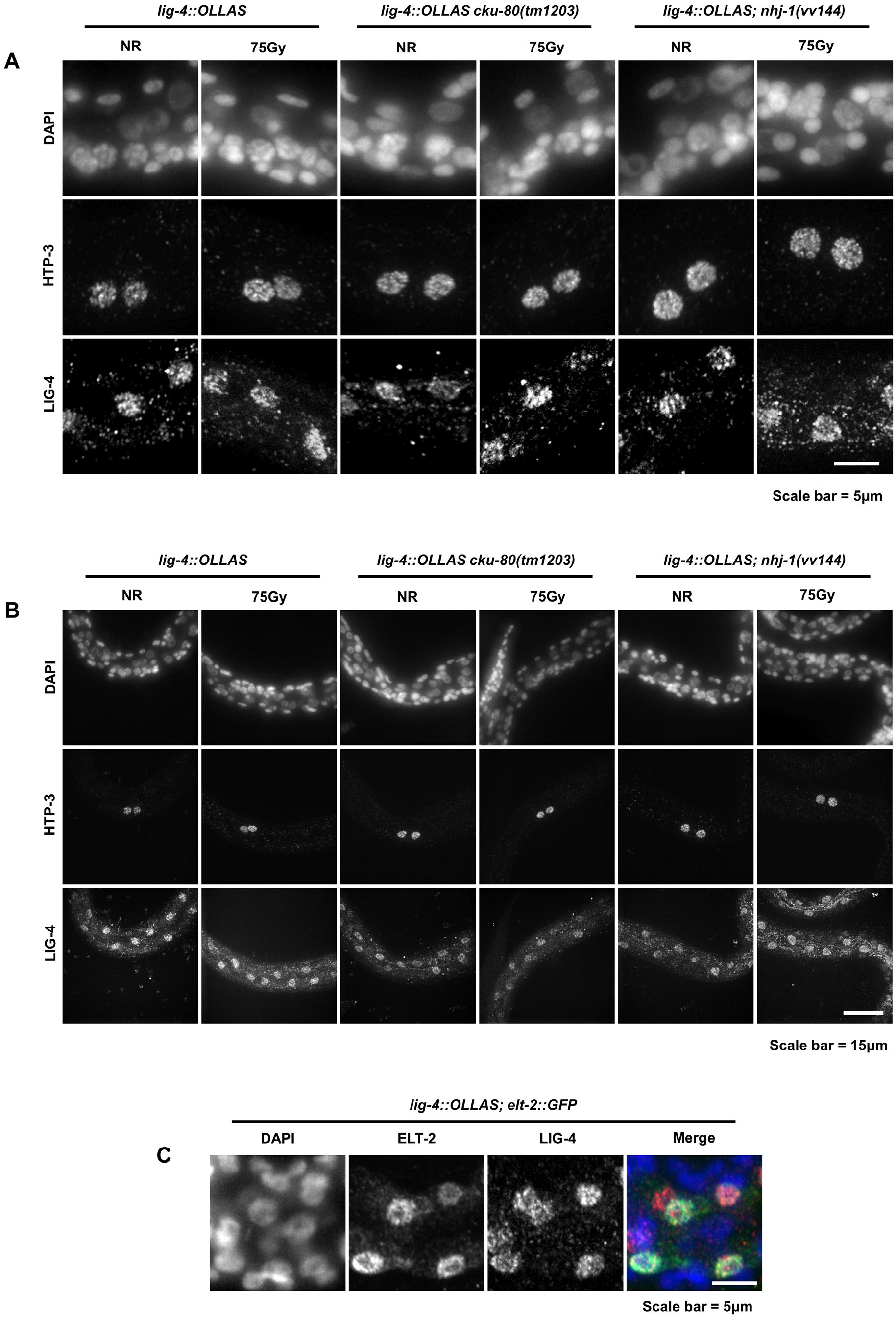
Endogenous LIG-4 localization in the L1. **(A)** Representative micrographs showing the subcellular localization of LIG-4::OLLAS from the endogenous locus, together with DNA staining (DAPI) and the germline marker HTP-3, in the L1 larva. The LIG-4 signal is detectable beyond background levels only in a row of nuclei along the anterior-posterior axis (see **(B)**). No LIG-4 signal is detected in the PGCs. **(B)** Representative micrographs showing the subcellular localization of LIG-4::OLLAS, HTP-3, and DNA in the same genotypes and conditions as in **(A)**, but in a wider field of view, showing the enrichment in a longitudinal row of nuclei. **(C)** Representative micrograph showing the nuclear co-localization of LIG-4::OLLAS and ELT-2::GFP, an intestinal cell marker. The nuclei which most strongly express LIG-4 also express the intestinal marker ELT-2::GFP, suggesting that LIG-4 is enriched in the intestine.

**Figure 12.**
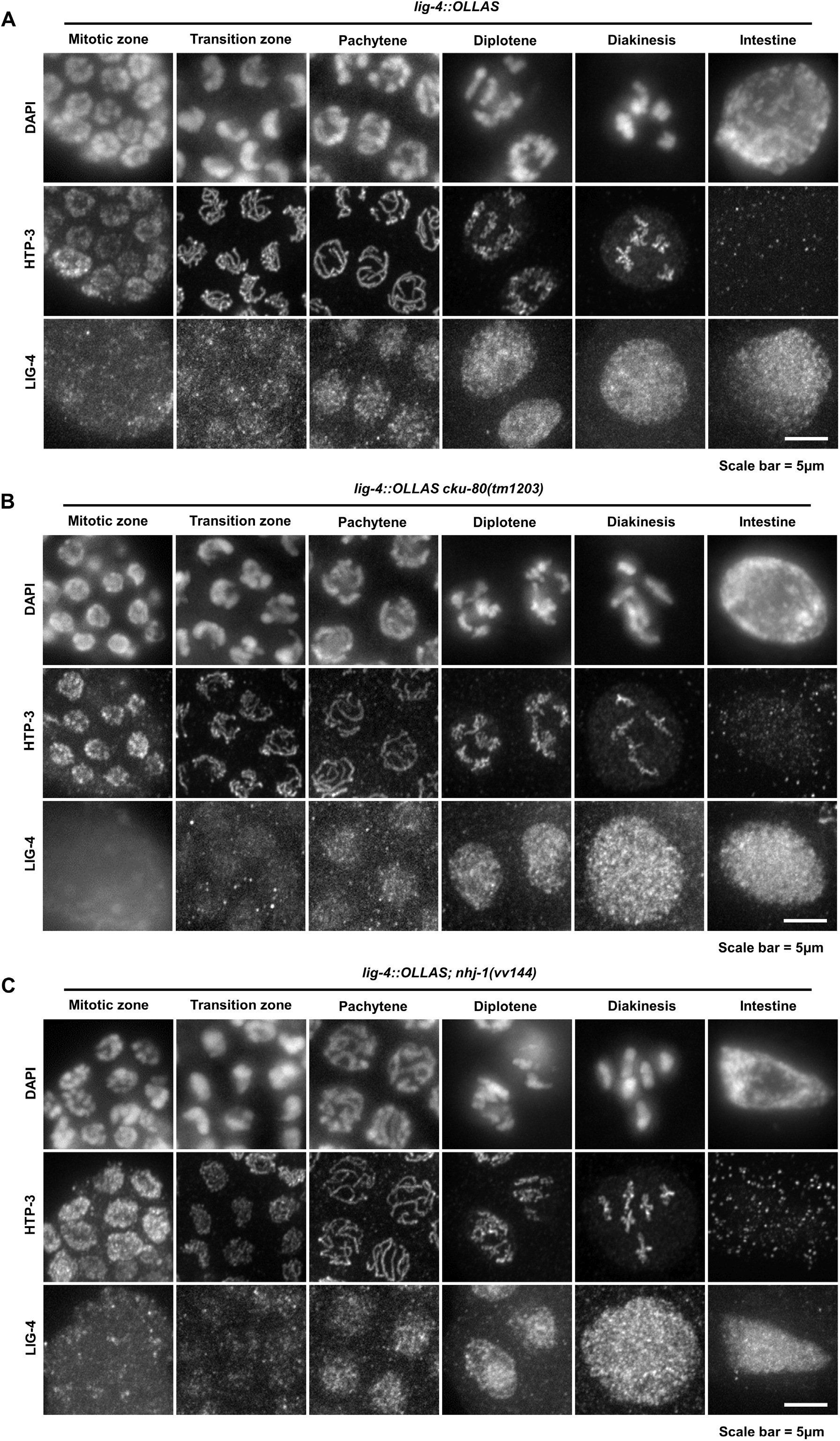
Endogenous LIG-4 localization in the adult germline. **(A)** Representative micrographs of LIG-4::OLLAS expression from the endogenous locus in adult germline cells. The expression of LIG-4::OLLAS becomes reliably visible in pachytene, and is nuclear, punctate, and not chromatin associated. An adult intestinal cell, where LIG-4 is also strongly expressed, is shown for comparison. **(B)** Representative micrographs of LIG-4::OLLAS expression from the endogenous locus in adult germline cells in animals deficient for *cku-80.* Similar to NHJ-1::OLLAS, the loss of CKU-80 does not alter the localization of LIG-4::OLLAS either in the germline or in the intestine. **(C)** Representative micrographs of LIG-4::OLLAS expression from the endogenous locus in adult germline cells in animals deficient for *nhj-1.* The absence of NHJ-1 does not affect the localization pattern of LIG-4.

## Discussion

### A laboratory N2 line carries a radiosensitizing mutation

Here, we have presented the discovery of a critical role in canonical non-homologous end-joining for *H19N07.3/nhj-1*, a gene which has been recently reported to play a role in bleomycin resistance [18]. We initially observed a strongly divergent phenotypic response to ionizing radiation in several lines of the N2 strain, in which the sensitive N2 [S] line displayed a markedly reduced brood size as well as slow growth and vulval dysgenesis phenotypes that the resistant N2 [R] line did not, which prompted us to more closely examine the phenomenon. Treatment with ethyl-nitrosourea and UV radiation strongly suggested that the sensitive line N2 [S] is specifically sensitive to IR, and not other genotoxic stressors, and only at the early larval stages. Since this IR-sensitive phenotype segregated in a Mendelian pattern, we sequenced the N2 [S] and N2 [R] genomes, and using homozygous variants as molecular markers, mapped a candidate for the causative variant to an indel in the *H19N07.3/nhj-1* locus. We concluded that the loss of *nhj-1* is responsible for IR sensitivity of N2 [S] since inactivation of *nhj-1* by CRISPR-Cas9 in the N2 [R] genome was sufficient to induce radiation sensitivity and the CRISPR allele *vv144* was unable to complement the natural allele *vv148*. Since unannotated cryptic genetic variation has been documented to occur in laboratory strains of *C. elegans* as a result of drift [29], this finding was surprising only in the magnitude of the effect.

### *H19N07.3/nhj-1 is a novel C. elegans* cNHEJ factor

Because IR efficiently induces DNA DSBs and a cNHEJ deficiency has been observed in *C. elegans* to cause somatic phenotypes similar to the ones we observed in N2 [S] [17], we investigated the possibility that N2 [S] is sensitive because of a loss of a cNHEJ factor. In genetic *nhj-1; lig-4* and *nhj-1; cku-80* double mutants, we did not observe additive IR sensitivity, suggesting that *nhj-1* is a cNHEJ factor. This conclusion is further supported by an increase in the number of DAPI bodies in *com-1; nhj-1* mutants, compared to *com-1* mutants alone, which is also observed in *com-1; lig-4* mutants, and is consistent with the interpretation that the increased DNA fragmentation in *com-1; nhj-1* mutants results from a loss of cNHEJ activity. The recent findings that NHJ-1 is required for resistance to bleomycin, a chemotherapeutic agent that can cause DNA DSBs, [18] accord with our results of a role in cNHEJ. With the notable exception of vertebrates, who rely on cNHEJ factors for V(D)J recombination to generate an adaptive immune response [30], cNHEJ is dispensable for survival in eukaryotes. This may have allowed an inactivating mutation in a critical cNHEJ factor to spread in laboratory populations.

*C. elegans* was thought to either possess a minimal cNHEJ system, composed of only the Ku ring and LIG-4, or that other cNHEJ components such as nucleases or kinases, had yet to be identified because no saturated screen for cNHEJ factors has been done [16]. While the role of *nhj-1* in the cNHEJ pathway remains opaque, the increased number of diakinetic DAPI bodies in *com-1; nhj-1* double mutants compared to *com-1* single mutants suggests that it acts downstream of Ku, and likely upstream of LIG-4, which performs the terminal ligation step. The roles NHJ-1 may play downstream of Ku binding include: 1) DNA end processing; 2) Signaling to coordinate the activity or assembly of the cNHEJ complex; 3) Promoting the activity of other cNHEJ pathway components as a cofactor; and 4) Acting as a structural scaffold to organize and coordinate other cNHEJ factors. Our analysis and that of others [18] shows that NHJ-1 contains no conserved domains. Together with the relatively small size of the protein (the longer of the two isoforms is 168 amino acids long), we consider the possibility that NHJ-1 is an enzyme unlikely. Signaling and processing enzymes with active roles in cNHEJ tend to be much larger, with the ∼4,000 amino acid-long DNA-PKcs at the higher end of the spectrum, and the ∼500 amino acid-long nuclease APLF at the lower end [1]. By contrast, the structural proteins XRCC4, XLF, and PAXX, are of much more modest size, ranging from 201 aa (PAXX) to 334 aa (XRCC4) in *H. sapiens* [1], although XRCC4 and XLF have been shown to oligomerize into much larger filaments that support other cNHEJ machinery [31, 32]. NHJ-1 could act in an analogous manner in *C. elegans*, even though it shares no sequence homology with XRCC4 or its homologs. However, a role in an enzyme-driven step of cNHEJ cannot be definitively excluded, as NHJ-1 could act as a co-factor to promote enzymatic activity or even possess enzymatic activity itself.

As previously noted [18], the scope of the evolutionary conservation of NHJ-1 is limited to closely related nematodes. Proteins with high identity with the NHJ-1 long isoform exist in several species of the genus *Caenorhabditis*, including *C. brenneri* (90% identity), *C. briggsae* (88% identity), *C. remanei* (84% identity), and *C. latens* (84% identity). In the family Rhabditidae, which includes the genus *Caenorhabditis,* there are two homologs in the asexual worm *Diploscapter pachys* (33% and 32% identity). The only other proteins with homology belong to two parasitic hookworms in the family Ancylostomatidae, *Necator americanus* (24% identity), and *Ancylostoma duodenale* (22% and 20% identity). What roles the homologs of NHJ-1 play in the other nematodes is not known. However, given the relatively high sequence conservation within *Caenorhabditis*, NHJ-1 homologs in the other species of this genus may also participate in cNHEJ. The nematode family Rhabditidae, thus appears to have evolved a novel regulator of the nearly universally conserved [12] cNHEJ pathway, illustrating the evolutionary plasticity of even the most ancient pathways. The lack of sequence conservation between NHJ-1 and the known cNHEJ factors in other phyla also highlights the possibility that the cNHEJ toolkit in *C. elegans* may be much larger, and novel functional analogs to other cNHEJ factors may yet be discovered.

### NHJ-1 localization is consistent with a role in cNHEJ

We also examined the subcellular localization of NHJ-1 and LIG-4 in L1 larvae and adult gonad and intestinal tissue, both of which raised interesting questions about the regulation of cNHEJ in specific tissue contexts. The localization of cNHEJ components, the regulation of their recruitment to sites of DNA damage, and their dependence on other cNHEJ factors for nuclear recruitment has primarily been studied in the context of cultured mammalian cells [33–37]. As expected for DNA repair factors, Ku, DNA-PKcs, XRCC4, XLF, PAXX, and LIG4 are predominantly nuclear, with Ku and possibly others excluded from the nucleus only during mitosis [33, 35, 36, 38–40]. By contrast, few published studies have examined the localization of cNHEJ factors in the tissue or organ context. In healthy human colon tissue, Ku70 is detectable by immunohistochemistry in 74% of nuclei, in contrast to Ku80, which can be seen in only 32% of nuclei [41]. Similarly, many but not all cells in the crypts of human and murine small intestine express LIG4, which is detectable in both nuclei and the cytoplasm in the cells that express it [42]. In the mouse testis, Ku70 localizes to the nuclei of the somatic Sertoli cells, spermatogonia, late (post-pachytene) spermatocytes I, spermatocytes II, and spermatids [43].

These observations are concordant with our localization data. In *C. elegans*, NHJ-1 and LIG-4 are nuclearly localized in both the L1 larva and the adult gonad and intestine. In the adult germline, the two proteins have a similar expression pattern, with strongest expression in diplotene and diakinesis, although LIG-4 becomes visible in pachytene. This is in line with the role of cNHEJ as a backup DNA repair pathway during meiotic prophase I, when inter-homolog HR repair is heavily favored [16, 17, 44, 45]. This expression pattern is also reminiscent of the localization of Ku70 in the mouse seminiferous tubules [43], except that some Ku70 expression is seen in spermatogonia while neither NHJ-1 nor LIG-4 are visible in the mitotic zone of *C. elegans*. The tissue-level expression pattern of NHJ-1 and LIG-4 in the L1 larva is markedly different, however. While NHJ-1 localizes to many somatic nuclei, LIG-4 is detectable primarily in the intestine. Several possibilities exist that could explain this discordance. NHJ-1 could be pleiotropic and possess a cNHEJ-independent function in non-intestinal cells, although it is unlikely that cNHEJ would operate only in the intestinal cells at the L1 stage. Expression levels of LIG-4 may be below the detection threshold in non-intestinal nuclei, which would suggest a relative enrichment of LIG-4 in intestinal cells compared to other tissues. Although intestinal cell nuclei are diploid in the early L1 larva, their ploidy doubles with each larval stage to the final number of 32 copies of each chromosome in adult [46], suggesting that the LIG-4 enrichment may reflect a greater need for cNHEJ in this tissue. In addition to increased ploidy, an increased requirement for cNHEJ in the intestinal cells may result from the fact that these cells are the ones most likely to be directly exposed to toxins produced by pathogenic bacteria and other microbiota which can colonize the intestinal lumen [47]. However, NHJ-1 is not enriched in intestinal nuclei compared to other somatic nuclei, suggesting that a general enrichment of cNHEJ factors is not sufficient to explain the LIG-4 pattern.

Our study has revealed *nhj-1* as a novel player in the *C. elegans* cNHEJ toolkit, and is in agreement with the recent findings that *nhj-1*/*scb-1* is required for resistance to bleomycin, a chemotherapeutic agent that induces DNA DSBs [18]. The challenge now is to elucidate the mechanistic role of NHJ-1 in the cNHEJ process, as well as to identify other potential cNHEJ factors in *C. elegans* and related nematodes.

## Materials and methods

### *Caenorhabditis* strain maintenance and mating

All *C. elegans* and *C. briggsae* strains have been maintained under standard conditions, at 20°C on Nematode Growth Medium (NGM) with the *E. coli* strain OP50 as a food source [48, 49]. The sensitive N2 [S] strain was derived from a single animal isolated from the N2 line from the *Caenorhaditis* Genetics Center. The IR-resistent N2 [R] strain was derived from a single animal isolated from the N2 line generously supplied by Dr. Erik Andersen. For a list of strains used in this study, see **Table S4**.

### Ionizing radiation treatment

Animals were treated with ionizing radiation in the form of X-rays (RS 2000 small animal X-ray irradiator, Rad Source Technologies Inc) at the rate of 2.34 Gray per minute. Control animals were kept next to the IR source during the irradiation.

For homozygous L1 animals, irradiation was performed in M9 buffer in 1.5 ml microcentrifuge tubes. Synchronized L1 animals were obtained by hypochlorite treatment as previously described [50], with the following modifications: washes were done in water instead of M9 buffer, the animals were treated with hypochlorite for 10-12 minutes, and the hypochlorite solution recipe used was 3.3 ml water, 1.2 ml sodium hypochlorite (4%), 0.5 sodium hydroxide (0.5M). After hypochlorite treatment, the eggs were left overnight in M9 buffer in 15 ml centrifuge tubes to hatch. Two hours before irradiation, a concentrated culture of OP50 *E. coli* in the amount totaling 10% of the M9 buffer volume was added to the 15 ml centrifuge tubes as a food source for the L1-arrested larvae. Immediately before irradiation, the larvae were transferred to 1.5 ml microcentrifuge tubes, and irradiated as described above. Following irradiation, the animals were transferred by glass Pasteur pipettes to fresh NGM plates.

For heterozygous and homozygous L1 cross-progeny, irradiation was performed on NGM plates. Near-synchronized cross-progeny L1s were obtained by isolating mated hermaphrodites on NGM plates, allowing them to lay eggs for 4 hours, and then removing them from the plates. The plates containing hatched L1 larvae were irradiated 12-14 hours after the removal of the mothers.

For L4 animals, irradiation was performed on NGM plates. Animals were synchronized using hypochlorite treatment as described above and dispensed onto NGM plates. After 48 hours, the L4 animals were irradiated directly on the plate.

### ENU treatment

For ENU treatment, synchronized L1 larvae were obtained by hypochlorite bleaching and provided with OP50 as described above. After allowing 2 hours for feeding, the larvae were incubated in a 15 ml centrifuge tube with the working solution of ENU as described in [51], except: L1 larvae were used instead of L4 larvae, and the working concentration of ENU used was 5 mM and 10 mM. Control animals were kept in tubes containing only M9, next to the ENU tubes.

### UV treatment

For UV irradiation, synchronized L1 larvae were similarly obtained by the hypochlorite bleaching method. Following synchronization, the L1 larvae were dispensed on NGM plates with a glass Pasteur pipette, and irradiated on plates with 50 J/m^2^ or 100J/m^2^ of UV-C in a Stratalinker 1800 UV crosslinker (Stratagene California). Control animals were kept on plates next to the crosslinker.

### Scoring of somatic phenotypes

Following IR, UV, or ENU exposure, animals were transferred onto NGM plates (or left on the plate if treated on plates), and left to develop for three days (72-76 hours) or four days (96-100 hours). At those time points, the incidence of somatic phenotypes was assessed in the following way. Protruding vulva, ruptured through vulva, and larvae were scored directly on the plate using a Leica MS5 stereomicroscope. A variant method was used in experiment scoring phenotypes in cross progeny. Here, the animals were isolated to individual plates as L1 following treatment, and the somatic phenotype scored for each animal at three and four days after treatment.

The proportion of worms showing the Egl phenotype was scored by dissecting individual animals using hypodermal injection needles (Becton, Dickinson and Company). Animals were scored as Egl if they contained one or more hatched larvae within their body at time of dissection (bagging).

### Scoring of embryonic lethality

Groups of IR-treated and control animals were moved to a fresh NGM plate 24 hours after the L4 stage, and moved to a fresh plate two times after that in 8 hour intervals. After 8 hours on the last plate, the animals were removed, leaving three brood plates. The number of eggs was scored on each plate following the transfer or removal of the animals, and the number of hatched larvae was counted on each plate ∼24 hours after the transfer or removal of the mothers. For L1 treated animals, 10-12 animals per plate were used for unirradiated and IR-treated resistant groups, and 20 animals per plate for IR-treated sensitive groups. For animals treated at L4, 10 animals per plate were used for unirradiated controls and 12-15 animals were used for IR-treated groups.

### CRISPR-Cas9 mutagenesis

CRISPR-Cas9 mutagenesis was performed using the *in vitro* assembled ribonucleoprotein complex as described [52]. Young adult animals were microinjected the Cas9/tracrRNA/crRNA/DNA repair template mixture 1 day after L4 stage. The *dpy-10(cn64)* allele was used as a co-conversion marker [53]. Heterozygous *dpy-10(cn64/+)* rollers were isolated from the progeny of injected animals, allowed to lay progeny, then lysed and genotyped for the mutation of interest. If positive, wild-type moving F2 progeny was isolated and genotyped for homozygosity of the mutation of interest. TracrRNA and crRNAs were synthesized by GE Healthcare Dharmacon, Inc. The repair template DNA oligonucleotides were synthesized by Integrated DNA Technologies, Inc. The Cas9 endonuclease was purchased from PNA Bio Inc. The crRNAs and repair templates used to create mutations used in this study are listed below.

*lig-4* N-term crRNA (DNA target): TTGACGTCTTCAACAAGATT

*lig-4(vv134*[*R18STOP*]) repair template:

ATGGCGTCAGATGTGATCTTCGACGAAGTAGTTGACGTCTTCAACAAGATTTGACGGACTTCAAATGTGAAATCAAAGCAAGCAACCTTTCAGAAAAACTTTGAATCATGGAA AG

*lig-4* C-term crRNA (DNA target): CGAAGGTGGATTCGAGATTC

*lig-4(vv145*[*lig-4::OLLAS*]) repair template:

TGGTTGCCTTCTGATGTGTTTCATGCCATCGAAGGTGGATTCGAGATTCAGGAATACCCATATGATGTCCCGGATTACGCTTAATTTACTAATTTCGATTATATGTGATATCGCTCTTTATTTCCTTTTT

*nhj-1* crRNA (DNA target): CTAGAGCGTACGGAGCTTTC

*nhj-1(144)* repair template:

TTCCCTCTTCTCTGAAAGTGGCCTTCATATCGAAGAAGTTTGTGAGAAGGAGCTTGTGCTCACGTTTGCACTTCAAAGAAATATTAGCGTACGGAGCTTTCTGGAAGATTCGATGAGTGCCTCTACTTGGAATATTTGGTGGGTTAAAAAGTT

*nhj-1* C-term crRNA (DNA target): ATCGTCAAACTCTGGTCCAC

*nhj-1(vv147*[*nhj-1::OLLAS*]) repair template:

TATGGCTGCCAAGGCCAGTGGACCAGAGTTTGACGATGAATCTGGATTCGCTAACGAGCTTGGACCACGCCTTATGGGAAAGTAATTTCACAATTAATTAACCCCATCTTTCTTGTTCCATG

### Genomic DNA preparation and sequencing

Genomic DNA for deep sequencing was prepared using the Schedl lab protocol (http://genetics.wustl.edu/tslab/protocols/genomic-stuff/worm-genomic-dna-prep). Briefly, 5 medium-sized (10 cm diameter) NGM plates were seeded with 15 L4 animals and allowed to starve over the course of a week. The arrested L1 larvae were then grown in liquid NGM 3-4 days until they developed into adults, which were treated with a 30% sucrose float to remove food contamination and separated into 500 μl aliquots, which were frozen at −80°C. An aliquot was then transferred to a 15 ml centrifuge tube, 4.5 ml of worm lysis buffer (0.1M Tris-Cl pH 8.5, 0.1M NaCl, 50 mM EDTA pH 8.0, 1% SDS) and 200 μl of Protease K (20 mg/ml in TE pH 8.0) added, and the worms vortexed. The mixture was incubated for 1 hour at 62°C, with intermittent vortexing. Then, 800 μL of 5M NaCl and was added and the tube mixed by inversion, after which 800 μL of CTAB solution (10 % CTAB in 0.7M NaCl) was added and the tube incubated for 10 minutes at 37°C. Following this, 7 ml of chloroform was added and the tube mixed and spun, the aqueous phase recovered, and the step repeated with 7 ml phenol/chloroform/isoamyl alcohol. Next, 0.6 volume of −20°C isopropyl alcohol was added, and mixed, and the DNA spun at 4°C for 5 minutes. The DNA pellet was washed in 70% ethanol, dried, and resuspended in 340 μl of TE buffer. Next, 10 μl of RNase A (10 mg/ml) was added and the tube incubated for 2 hours at 42°C, following which 20 μl of 20 % SDS, 10 μl of 0.5 M EDTA pH 8.0, and 20 μl of Protease K was added and the tube incubated for 2 hours at 65°C. Then, 40 μl of 10 M Ammonium Acetate was added, the DNA extracted twice with phenol/chloroform/isoamyl alcohol and once with chloroform, 1 ml of ethanol added, and the DNA spun down at 4°C for 10 minutes. The DNA was washed twice with 70% ethanol, dried, and resuspended in 200 μl of TE buffer.

Sample paired-end tag libraries were prepared by Canada’s Michael Smith Genome Sciences Centre, and the samples were sequenced with Illumina HiSeq 2500 (125 bp read length) to a coverage of 100X for N2 [S] and 200X for N2 [R]. The Genome Sciences Centre also provided the binary alignment (bam) files for both genomes.

### Bioinformatic analysis

Sequence variants in N2 [S] and N2 [R] were called with SAMtools [54], using the mpileup function against the WS249_cel235.fa reference genome. Filtering was performed and strain-specific variants determined using the somatic variation function in the small variant caller Strelka2 [55], except the *nhj-1(vv148)* mutation, which was identified by manual parsing through variants called by SAMtools mpileup. The full sequence of the *nhj-1(vv148)* indel was identified by N2 [S] genome reassembly with ABySS 2.0 [56] from the sorted bam file.

The search for protein sequences homologous to NHJ-1 was conducted with DELTA-BLAST on the NCBI online tool, https://blast.ncbi.nlm.nih.gov/Blast.cgi [57]. NHJ-1 isoform sequences were analyzed for domain conservation by SMART (Simple Molecular Architecture Research Tool), http://smart.embl-heidelberg.de [58]. The hairpin in *nhj-1(vv148)* insertion was predicted using the ViennaRNA package 2.0 [59].

### Mapping

The mapping of the IR-sensitivity-causative locus (*nhj-1*) was done in two rounds, using N2 [S]- and N2 [R]-specific molecular markers identified through deep sequencing to assay marker segregation in F2 hybrid strains originating from an N2 [S] X N2 [R] cross. In the first, low-resolution mapping round, N2 [S]-specific candidate loci were PCR amplified and Sanger sequenced at the Genome Quebec center at McGill University Campus. In the second, high-resolution mapping round, the variants in *F10D2.12* and *inft-2* were used as RFLPs, with KpnI cutting the [R]-form of *F10D2.12* but not the [S]-form and TfiI cutting the [R]-form of *inft-2*, but not the [S]-form.

### Genotyping of the *nhj-1* locus

The wild type *nhj-1* locus and *nhj-1(vv144)* were amplified using the following gene flanking primers:

F: TTGTGTTGAAACTGTACCGTCT; and

R: CAAAGTAGTCCCCCTAATCGCA.

Digestion of the resulting product with XbaI yields two bands on *nhj-1(+)* but does not cut *nhj-1(vv144)*. The flanking primer pair does not yield a product with *nhj-1(vv148)*, likely because of the hairpin present in this allele, and *nhj-1(vv148)* was therefore genotyped using an alternate reverse primer which anneals in the hairpin section:

F: TTGTGTTGAAACTGTACCGTCT; and

R: TAATAATATTTTTAATAAATAATAGTAATAT.

### Immunostaining

Gonad and intestinal immunohistochemistry was performed on dissected organs as previously described [60], with the following adjustments: dissection was performed in M9 buffer, four washes with PBST preceded the blocking, blocking was done with 1% BSA in PBST, slides were incubated with primary antibodies at 4°C, slides were washed with 1% BSA PBST four times before addition of secondary antibodies, incubated with secondary antibodies for 2 hours at room temperature, and washed again four times with PBST before addition of DAPI in Vectashield mounting medium (Vector Laboratories Inc).

Immunohistochemistry of L1 larvae was performed using the freeze-crack method as described in [61], with the following modifications: slides were left in −20°C for 1 minute, fixed with 1% formaldehyde in PBST for 5 minutes, washed four times with PBST before blocking with 1% BSA in PBST, washed four times with 1% BSA in PBST before secondary antibody incubation, and washed four times with PBST before addition of DAPI in Vectashield.

The primary antibodies and concentrations used in this study are as follows: guinea pig α-HTP-3 (1:500) [62], rat α-OLLAS (1:200) (Novus Biologicals, Inc), mouse α-GFP (1:200) (Abcam), rabbit α-H3K9me3 (1:500) (Cell Signaling Technology, Inc), and rabbit α-H3K9Ac (1:200) (Cell Signaling Technology, Inc).

Secondary antibodies used in this study include: Alexa 488-conjugated α-guinea pig, Alexa 488-conjugated α-mouse, Alexa 555-conjugated α-rabbit, and Alexa 555-conjugated α-rat. All secondary antibodies were purchased from Molecular Probes Inc, and used at a concentration of 1:1000.

### Microscopy

All worm manipulations, transfers, and crosses, as well as brood size scoring, somatic phenotype scoring, embryonic lethality scoring, and worm dissection was performed on Leica MS5 stereomicroscopes.

Example somatic phenotypes shown in Figure 2 were imaged with a 12-bit QICAM digital camera (QImaging and Photometrics) on a Leica MZ8 stereomicroscope.

Micrographs shown in Figures 8-12 were acquired with a Leica DMI 6000B inverted microscope and EM CCD camera C1900 (Hamamatsu Photonics KK). The DAPI signal was acquired with wide-field X-Cite 120 florescence illumination system (Excelitas Technologies), while the Alexa-488 and Alexa-555 conjugated antibody signals were acquired with a Quorum WaveFX spinning disc confocal system (Quorum Technologies), both integrated with the Leica DMI 6000B microscope. Images were acquired in stacks of 15-40 Z-planes in increments of 0.2 μm. Stack projections and contrast and brightness adjustments were performed in ImageJ (National Institutes of Health and Laboratory for Optical and Computational Instrumentation).

### RNAi of *cku-80*

RNAi knockdown of *cku-80* was done according to the standard feeding protocol [63]. Heterozygous *com-1* animals were put on plates containing *cku-80* expressing bacteria at the L4 stage, and the F1 progeny individually plated on *cku-80* RNAi plates for scoring of embryonic lethality. Control animals were fed bacteria expressing the empty vector L4440.

### Statistical analyses, descriptive statistics, and data presentation

Because of the non-normal distribution observed in the post-IR brood size data, this data was compared with the non-parametric Kruskal-Wallis H-test [64]. Pairwise comparisons between individual groups were done by Dunn’s post-hoc test [65] or serial Mann-Whitney U-tests [66] with a Bonferroni correction [67] applied to compensate for multiple testing. Categorical data, including the incidence of post-IR somatic phenotypes and embryonic lethality, is analyzed by Pearson’s Chi-squared test [68], and in cases of multiple comparisons compared against a Bonferroni-corrected α-value.

Because of the non-normal distribution of the brood size data and the non-parametric tests used to determine significance, the descriptive statistical metrics used both as error bars in the figures and reported in the text are the median and the interquartile range, rather than the mean and standard deviation.

All statistical tests were performed in GraphPad Prism 5 (GraphPad Software Inc). Vertical scatter plots were generated in GraphPad Prism 5, and 100% stacked column bar graphs were generated in Microsoft Excel (Microsoft Corporation).

## Acknowledgements

This work was supported by funding from the Canadian Institute of Health Research (201109) and the Natural Sciences and Engineering Council (RGPIN-2018-05963) to MZ. We would like to thank Florence Couteau and members of the Zetka laboratory for helpful discussions and Sara Labella for experimental support. We are grateful to Anja Bošković, Maia Kaplan, and Harwood Kwan for bioinformatic data analysis. Some strains were provided by the CGC, which is funded by the NIH Office of Research Infrastructure Programs (P40 OD010440). We would also like to thank Erik Andersen, Siegfried Hekimi, Mei Zhen, and the National Bioresource Project (Japan) for strains, and the Roy Laboratory for strains, reagents, and critical discussions during the course of this work.

## References

1. Chang, H.H.Y., Pannunzio, N.R., Adachi, N., and Lieber, M.R. (2017). Non-homologous DNA end joining and alternative pathways to double-strand break repair. Nat Rev Mol Cell Biol 18, 495–506.

2. Betermier, M., Bertrand, P., and Lopez, B.S. (2014). Is non-homologous end-joining really an inherently error-prone process? PLoS Genet 10, e1004086.

3. Ceccaldi, R., Rondinelli, B., and D’Andrea, A.D. (2016). Repair Pathway Choices and Consequences at the Double-Strand Break. Trends Cell Biol 26, 52–64.

4. Li, J., and Xu, X.Z. (2016). DNA double-strand break repair: a tale of pathway choices. Acta Bioch Bioph Sin 48, 641–646.

5. Decottignies, A. (2013). Alternative end-joining mechanisms: a historical perspective. Front Genet 4, 48.

6. Sallmyr, A., and Tomkinson, A.E. (2018). Repair of DNA double-strand breaks by mammalian alternative end-joining pathways. Journal of Biological Chemistry 293, 10536–10546.

7. Iliakis, G., Murmann, T., and Soni, A. (2015). Alternative end-joining repair pathways are the ultimate backup for abrogated classical non-homologous end-joining and homologous recombination repair: Implications for the formation of chromosome translocations. Mutat Res-Gen Tox En 793, 166–175.

8. Wang, C., and Lees-Miller, S.P. (2013). Detection and repair of ionizing radiation-induced DNA double strand breaks: new developments in nonhomologous end joining. Int J Radiat Oncol Biol Phys 86, 440–449.

9. Davis, A.J., and Chen, D.J. (2013). DNA double strand break repair via non-homologous end-joining. Transl Cancer Res 2, 130–143.

10. Radhakrishnan, S.K., Jette, N., and Lees-Miller, S.P. (2014). Non-homologous end joining: Emerging themes and unanswered questions. DNA Repair 17, 2–8.

11. Lieber, M.R. (2008). The mechanism of human nonhomologous DNA end joining. Journal of Biological Chemistry 283, 1–5.

12. Gu, J., and Lieber, M.R. (2008). Mechanistic flexibility as a conserved theme across 3 billion years of nonhomologous DNA end-joining. Genes Dev 22, 411–415.

13. Manova, V., and Gruszka, D. (2015). DNA damage and repair in plants - from models to crops. Frontiers in Plant Science 6.

14. Daley, J.M., Palmbos, P.L., Wu, D., and Wilson, T.E. (2005). Nonhomologous end joining in yeast. Annu Rev Genet 39, 431–451.

15. Sekelsky, J. (2017). DNA Repair in Drosophila: Mutagens, Models, and Missing Genes. Genetics 205, 471–490.

16. Lemmens, B.B.L.G., and Tijsterman, M. (2011). DNA double-strand break repair in Caenorhabditis elegans. Chromosoma 120, 1–21.

17. Clejan, I., Boerckel, J., and Ahmed, S. (2006). Developmental modulation of nonhomologous end joining in Caenorhabditis elegans. Genetics 173, 1301–1317.

18. Brady, S.C., Zdraljevic, S., Bisaga, K.W., Tanny, R.E., Cook, D.E., Lee, D., Wang, Y., and Andersen, E.C. (2019). A Novel Gene Underlies Bleomycin-Response Variation in Caenorhabditis elegans. Genetics 212, 1453–1468.

19. Povirk, L.F. (1996). DNA damage and mutagenesis by radiomimetic DNA-cleaving agents: bleomycin, neocarzinostatin and other enediynes. Mutat Res 355, 71–89.

20. Barriere, A., and Felix, M.A. (2005). High local genetic diversity and low outcrossing rate in Caenorhabditis elegans natural populations. Curr Biol 15, 1176–1184.

21. Sulston, J.E., and Horvitz, H.R. (1977). Post-embryonic cell lineages of the nematode, Caenorhabditis elegans. Dev Biol 56, 110–156.

22. Acevedo-Arozena, A., Wells, S., Potter, P., Kelly, M., Cox, R.D., and Brown, S.D. (2008). ENU mutagenesis, a way forward to understand gene function. Annu Rev Genomics Hum Genet 9, 49–69.

23. Rastogi, R.P., Richa, Kumar, A., Tyagi, M.B., and Sinha, R.P. (2010). Molecular Mechanisms of Ultraviolet Radiation-Induced DNA Damage and Repair. J Nucleic Acids.

24. Sarov, M., Murray, J.I., Schanze, K., Pozniakovski, A., Niu, W., Angermann, K., Hasse, S., Rupprecht, M., Vinis, E., Tinney, M., et al. (2012). A genome-scale resource for in vivo tag-based protein function exploration in C. elegans. Cell 150, 855–866.

25. Lemmens, B.B.L.G., Johnson, N.M., and Tijsterman, M. (2013). COM-1 Promotes Homologous Recombination during Caenorhabditis elegans Meiosis by Antagonizing Ku-Mediated Non-Homologous End Joining. Plos Genetics 9.

26. Hunter, N. (2015). Meiotic Recombination: The Essence of Heredity. Cold Spring Harb Perspect Biol 7.

27. Park, S.H., Cheong, C., Idoyaga, J., Kim, J.Y., Choi, J.H., Do, Y., Lee, H., Jo, J.H., Oh, Y.S., Im, W., et al. (2008). Generation and application of new rat monoclonal antibodies against synthetic FLAG and OLLAS tags for improved immunodetection. J Immunol Methods 331, 27–38.

28. Francis, D.B., Kozlov, M., Chavez, J., Chu, J., Malu, S., Hanna, M., and Cortes, P. (2014). DNA Ligase IV regulates XRCC4 nuclear localization. DNA Repair (Amst) 21, 36–42.

29. Flibotte, S., Edgley, M.L., Chaudhry, I., Taylor, J., Neil, S.E., Rogula, A., Zapf, R., Hirst, M., Butterfield, Y., Jones, S.J., et al. (2010). Whole-genome profiling of mutagenesis in Caenorhabditis elegans. Genetics 185, 431–441.

30. Woodbine, L., Gennery, A.R., and Jeggo, P.A. (2014). The clinical impact of deficiency in DNA non-homologous end-joining. DNA Repair (Amst) 16, 84–96.

31. Hammel, M., Rey, M., Yu, Y., Mani, R.S., Classen, S., Liu, M., Pique, M.E., Fang, S., Mahaney, B.L., Weinfeld, M., et al. (2011). XRCC4 protein interactions with XRCC4-like factor (XLF) create an extended grooved scaffold for DNA ligation and double strand break repair. J Biol Chem 286, 32638–32650.

32. Mahaney, B.L., Hammel, M., Meek, K., Tainer, J.A., and Lees-Miller, S.P. (2013). XRCC4 and XLF form long helical protein filaments suitable for DNA end protection and alignment to facilitate DNA double strand break repair. Biochem Cell Biol 91, 31–41.

33. Koike, M., Shiomi, T., and Koike, A. (2001). Dimerization and nuclear localization of ku proteins. J Biol Chem 276, 11167–11173.

34. Koike, M., Awaji, T., Kataoka, M., Tsujimoto, G., Kartasova, T., Koike, A., and Shiomi, T. (1999). Differential subcellular localization of DNA-dependent protein kinase components Ku and DNA-PKcs during mitosis. J Cell Sci 112, 4031–4039.

35. Nilsson, A., Sirzen, F., Lewensohn, R., Wang, N., and Skog, S. (1999). Cell cycle-dependent regulation of the DNA-dependent protein kinase. Cell Prolif 32, 239–248.

36. Girard, P.M., Kysela, B., Harer, C.J., Doherty, A.J., and Jeggo, P.A. (2004). Analysis of DNA ligase IV mutations found in LIG4 syndrome patients: the impact of two linked polymorphisms. Hum Mol Genet 13, 2369–2376.

37. Mari, P.O., Florea, B.I., Persengiev, S.P., Verkaik, N.S., Brueggenwirth, H.T., Modesti, M., Giglia-Mari, G., Bezstarosti, K., Demmers, J.A.A., Luider, T.M., et al. (2006). Dynamic assembly of end-joining complexes requires interaction between Ku70/80 and XRCC4. P Natl Acad Sci USA 103, 18597–18602.

38. Koike, M., Yutoku, Y., and Koike, A. (2015). Dynamic changes in subcellular localization of cattle XLF during cell cycle, and focus formation of cattle XLF at DNA damage sites immediately after irradiation. J Vet Med Sci 77, 1109–1114.

39. Ochi, T., Blackford, A.N., Coates, J., Jhujh, S., Mehmood, S., Tamura, N., Travers, J., Wu, Q., Draviam, V.M., Robinson, C.V., et al. (2015). DNA repair. PAXX, a paralog of XRCC4 and XLF, interacts with Ku to promote DNA double-strand break repair. Science 347, 185–188.

40. Yurchenko, V., Xue, Z., and Sadofsky, M.J. (2006). SUMO modification of human XRCC4 regulates its localization and function in DNA double-strand break repair. Mol Cell Biol 26, 1786–1794.

41. Mazzarelli, P., Parrella, P., Seripa, D., Signori, E., Perrone, G., Rabitti, C., Borzomati, D., Gabbrielli, A., Matera, M.G., Gravina, C., et al. (2005). DNA end binding activity and Ku70/80 heterodimer expression in human colorectal tumor. World J Gastroenterol 11, 6694–6700.

42. Jun, S., Jung, Y.S., Suh, H.N., Wang, W., Kim, M.J., Oh, Y.S., Lien, E.M., Shen, X., Matsumoto, Y., McCrea, P.D., et al. (2016). LIG4 mediates Wnt signalling-induced radioresistance. Nat Commun 7, 10994.

43. Ahmed, E.A., Sfeir, A., Takai, H., and Scherthan, H. (2013). Ku70 and non-homologous end joining protect testicular cells from DNA damage. J Cell Sci 126, 3095–3104.

44. Adamo, A., Collis, S.J., Adelman, C.A., Silva, N., Horejsi, Z., Ward, J.D., Martinez-Perez, E., Boulton, S.J., and La Volpe, A. (2010). Preventing nonhomologous end joining suppresses DNA repair defects of Fanconi anemia. Mol Cell 39, 25–35.

45. Smolikov, S., Eizinger, A., Hurlburt, A., Rogers, E., Villeneuve, A.M., and Colaiacovo, M.P. (2007). Synapsis-Defective mutants reveal a correlation between chromosome conformation and the mode of double-strand break repair during Caenorhabditis elegans meiosis. Genetics 176, 2027–2033.

46. Hedgecock, E.M., and White, J.G. (1985). Polyploid tissues in the nematode Caenorhabditis elegans. Dev Biol 107, 128–133.

47. Jiang, H., and Wang, D. (2018). The Microbial Zoo in the C. elegans Intestine: Bacteria, Fungi and Viruses. Viruses 10.

48. Brenner, S. (1974). The genetics of Caenorhabditis elegans. Genetics 77, 71–94.

49. Stiernagle, T. (2006). Maintenance of C. elegans. WormBook, 1–11.

50. Porta-de-la-Riva, M., Fontrodona, L., Villanueva, A., and Ceron, J. (2012). Basic Caenorhabditis elegans methods: synchronization and observation. J Vis Exp, e4019.

51. Kutscher, L.M., and Shaham, S. (2014). Forward and reverse mutagenesis in C. elegans. WormBook, 1–26.

52. Paix, A., Folkmann, A., Rasoloson, D., and Seydoux, G. (2015). High Efficiency, Homology-Directed Genome Editing in Caenorhabditis elegans Using CRISPR-Cas9 Ribonucleoprotein Complexes. Genetics 201, 47–+.

53. Arribere, J.A., Bell, R.T., Fu, B.X., Artiles, K.L., Hartman, P.S., and Fire, A.Z. (2014). Efficient marker-free recovery of custom genetic modifications with CRISPR/Cas9 in Caenorhabditis elegans. Genetics 198, 837–846.

54. Li, H., Handsaker, B., Wysoker, A., Fennell, T., Ruan, J., Homer, N., Marth, G., Abecasis, G., Durbin, R., and Genome Project Data Processing, S. (2009). The Sequence Alignment/Map format and SAMtools. Bioinformatics 25, 2078–2079.

55. Kim, S., Scheffler, K., Halpern, A.L., Bekritsky, M.A., Noh, E., Kallberg, M., Chen, X., Kim, Y., Beyter, D., Krusche, P., et al. (2018). Strelka2: fast and accurate calling of germline and somatic variants. Nat Methods 15, 591–594.

56. Jackman, S.D., Vandervalk, B.P., Mohamadi, H., Chu, J., Yeo, S., Hammond, S.A., Jahesh, G., Khan, H., Coombe, L., Warren, R.L., et al. (2017). ABySS 2.0: resource-efficient assembly of large genomes using a Bloom filter. Genome Res 27, 768–777.

57. Boratyn, G.M., Schaffer, A.A., Agarwala, R., Altschul, S.F., Lipman, D.J., and Madden, T.L. (2012). Domain enhanced lookup time accelerated BLAST. Biol Direct 7, 12.

58. Letunic, I., and Bork, P. (2018). 20 years of the SMART protein domain annotation resource. Nucleic Acids Res 46, D493–D496.

59. Lorenz, R., Bernhart, S.H., Honer Zu Siederdissen, C., Tafer, H., Flamm, C., Stadler, P.F., and Hofacker, I.L. (2011). ViennaRNA Package 2.0. Algorithms Mol Biol 6, 26.

60. Martinez-Perez, E., and Villeneuve, A.M. (2005). HTP-1-dependent constraints coordinate homolog pairing and synapsis and promote chiasma formation during C. elegans meiosis. Genes Dev 19, 2727–2743.

61. Butuci, M., Williams, A.B., Wong, M.M., Kramer, B., and Michael, W.M. (2015). Zygotic Genome Activation Triggers Chromosome Damage and Checkpoint Signaling in C. elegans Primordial Germ Cells. Dev Cell 34, 85–95.

62. Goodyer, W., Kaitna, S., Couteau, F., Ward, J.D., Boulton, S.J., and Zetka, M. (2008). HTP-3 links DSB formation with homolog pairing and crossing over during C. elegans meiosis. Dev Cell 14, 263–274.

63. Conte, D., Jr., MacNeil, L.T., Walhout, A.J., and Mello, C.C. (2015). RNA Interference in Caenorhabditis elegans. Curr Protoc Mol Biol 109, 26 23 21–30.

64. Kruskal, W.H., and Wallis, W.A. (1952). Use of Ranks in One-Criterion Variance Analysis. J Am Stat Assoc 47, 583–621.

65. Dunn, O.J. (1964). Multiple Comparisons Using Rank Sums. Technometrics 6, 241–&.

66. Mann, H.B., and Whitney, D.R. (1947). On a Test of Whether One of 2 Random Variables Is Stochastically Larger Than the Other. Ann Math Stat 18, 50–60.

67. Shaffer, J.P. (1995). Multiple Hypothesis-Testing. Annu Rev Psychol 46, 561–584.

68. Pearson, K. (1900). On the Criterion that a given System of Deviations from the Probable in the Case of a Correlated System of Variables is such that it can be reasonably supposed to have arisen from Random Sampling. Philos Mag 50, 157–175.

