## Supplemental Figures and Tables for "NHJ-1 regulates canonical non-homologous end joining in *Caenorhabditis elegans*"

### Supplementary Figures and Tables

**Figure S1. CGC N2 is IR-sensitive, while the Roy lab N2 is IR-resistant**

Quantification of brood size in N2 lines from the Zetka lab, the Roy lab, and the CGC. Following IR treatment at the L1 stage, the N2 line from the Caenorhabditis Genetics Center exhibits the same reduction in brood size as the N2 line from the Zetka lab ($p>0.05$). However, the N2 from the neighbouring laboratory of Dr. Richard Roy at McGill University is much more resistant, exhibiting a significantly higher brood size than either the Zetka lab N2 or the CGC N2 ($p<0.001$ for both comparisons), although still lower than in unirradiated controls ($p<0.01$). All statistical comparisons shown in the figure are to N2 (Zetka) 75 Gy (Kruskal-Wallis test and Dunn's post-hoc tests). Error bars represent the median and interquartile range. Sample size (n) is 40 for both N2 (Zetka) groups, 42 and 50 respectively for unirradiated and irradiated N2 (Roy), and 40 for both N2 (CGC) groups.

IR = ionizing radiation

Gy = Gray (unit)

CGC = Caenorhabditis Genetics Center

N2 (Zetka) = IR-sensitive N2 strain originating from the laboratory of Monique Zetka (McGill University)

N2 (Roy) = IR-resistant N2 strain originating from the laboratory of Richard Roy (McGill University)

ns = not significant ($p>0.05$); *** = $p<0.001$

**Figure S2. Mapping of the IR-sensitivity-causative locus**

**(A)** This pipeline shows the first line strategy in mapping. Twenty-one phenotypically IR-sensitive lines were derived from an N2 [S] X N2 [R] cross as shown here, and the 15 N [S]-specific, protein sequence-altering single nucleotide mutations sequenced in the hybrid IR-sensitive lines (**Figure 2.12**).

**(B)** For higher resolution mapping, another 75 phenotypically sensitive hybrid lines were derived, and *F10D2.12* and *inft-2* genotyped to identify recombination events between the causative locus and these two markers.

**(C)** This panel shows the possible recombination outcomes that explain the observed genotyping outcomes. The causative locus recombined away from *F10D2.12* in 4/75 strains and from *inft-2* in 1/75 strains, showing that it is located closer to *inft-2.*

FDR = false discovery rate; X = number of hybrid lines sequenced for each gene

**Figure S3. Loss of cNHEJ activity sensitizes the N2 [R] background to IR**

**(A)** Total brood size quantification of N2 [R], N2 [S], and the Ligase IV mutants *lig-4(ok716)* and *lig-4(vv134)* after IR treatment at the L1 stage. Both the published *lig-4(ok716)* deletion mutant (whose genetic background is unknown) and *lig-4(vv134)*, a null mutant the generated by CRISPR in the N2 [R] genetic background, exhibit a post-IR brood size not significantly different than N2 [S] or each other ($p>0.05$ for all comparisons). All three of these genotypes display a significantly reduced post-IR brood size compared to N2 [R] ($p<0.001$ for all comparisons to post-IR N2 [R]). All statistical comparisons shown in the figure are to irradiated *lig-4(vv134)* (Kruskal-Wallis test, followed by Dunn's post-hoc test). Error bars represent the median and interquartile range. Sample size (n) is 18 for unirradiated N2 [R], 58 for irradiated N2 [R], 20 for unirradiated N2 [S], 57 for irradiated N2 [S], 18 for unirradiated *lig-4(ok716)*, 56 for irradiated *lig-4(ok716)*, 25 for unirradiated *lig-4(vv134)*, and 50 for irradiated *lig-4(vv134).*

**(B)** Quantification of post-IR somatic phenotypes four days after IR treatment in the same groups as in **(A)**. N2 [S], *lig-4(ok716)*, and *lig-4(vv134)* animals show a much higher incidence of vulval phenotypes post-IR than do N2 [R] animals ($p<0.001$), while they are not significantly different than each other ($p>0.05$). All statistical comparisons shown in the figure are to *lig-4(vv134)* (Chi-squared test, Bonferroni corrected for multiple comparisons to $\alpha = 0.008$). Sample size (n) is 18 for unirradiated N2 [R], 58 for irradiated N2 [R], 20 for unirradiated N2 [S], 57 for irradiated N2 [S], 18 for unirradiated *lig-4(ok716)*,

56 for irradiated *lig-4(ok716)*, 25 for unirradiated *lig-4(vv134)*, and 50 for irradiated *lig-4(vv134).*

Pvl = protruding vulva phenotype

Rup = ruptured through vulva phenotype

"Thin" = thin, whitish L3-like larva

L4 = larva of the L4 stage

N2 [S] = sensitive N2 strain, derived from the CGC N2

N2 [R] = resistant N2 strain, derived from Andersen lab N2

ns = not significant ($p>0.05$ in **(A)**; $p>0.008$ in **(B)**), *** = $p<0.001$

**Table S1. List of wild type *Caenorhabditis* strains tested for L1 IR-sensitivity**

Seven N2 lines from different sources, 18 non-N2 *C. elegans* wild type isolates from all continents and diverse geographic latitudes, and two *C. briggsae* wild type isolates have been tested for the L1 response to IR. Only the N2s from the Zetka lab and the CGC are sensitive.

Zetka laboratory = laboratory of Dr. Monique Zetka, McGill University, Montreal, Canada

CGC = Caenorhabditis Genetics Center, University of Minnesota Twin Cities, United States of America

Roy laboratory = laboratory of Dr. Richard Roy, McGill University, Montreal, Canada

Hekimi laboratory = laboratory of Dr. Siegfried Hekimi, McGill University, Montreal, Canada

Andersen laboratory = laboratory of Dr. Eric Andersen, Northwestern University, Chicago, United States of America

Zhen laboratory = laboratory of Dr. Mei Zhen, Lunenfeld-Tanenbaum Research Institute, Toronto, Canada

NBRP = National Bioresource Project, Shinjuku-ku, Tokyo, Japan

IR = ionizing radiation

**Table S2. Embryonic lethality is increased in N2 [R] after IR treatment at the L1 stage, and in both N2 [R] and N2 [S] after IR treatment at the L4 stage**

The progeny of N2 [S] mothers irradiated with 75 Gy at the L1 stage do not exhibit an increased embryonic lethality compared to the progeny of unirradiated mothers, strongly suggesting that the reduced brood size of IR-treated N2 [S] animals is not caused by impaired or mutagenic DNA repair in the germline. The progeny of N2 [R] mothers irradiated with 75 Gy show a significantly higher embryonic lethality than either the progeny of unirradiated N2 [R] mothers or the progeny of N2 [S] mothers irradiated at the same dose ($p<0.001$ for both comparisons). Irradiation with 75 Gy at the L4 stage induces much higher embryonic lethality in the progeny in both N2 genetic backgrounds, in comparison to unirradiated controls ($p<0.001$ for both N2 [R] and N2 [S]). However, there is no significant difference in the level of embryonic lethality among the progeny of irradiated N2 [R] and N2 [S]. All statistical comparisons shown in the figure are to N2 [S] at the equivalent IR dose (Chi-squared test, Bonferroni corrected for multiple comparisons to $\alpha = 0.01$).

IR = ionizing radiation
Gy = Gray (unit)
N2 [S] = sensitive N2 strain, derived from the CGC N2
N2 [R] = resistant N2 strain, derived from Andersen lab N2
ns = not significant ($p>0.01$); *** = $p<0.001$

**Table S3. Unique variants which affect protein coding genes in N2 [S]**

This table presents the list of 15 fixed (homozygous) single nucleotide variants unique to N2 [S] which affect the protein sequence of the genes in which they are found. Thirteen variants result in amino acid substitutions, two introduce a premature stop codon, and one results in the abolition of a splice site. None of the variants affects a gene with a known role in DNA repair or DDR.

**Table S4. List of nematode strains used in this study**

This table lists all of the *Caenorhabditis* strains used in this study, including both those received from outside sources and created for the purposes of this study. All strains are *C. elegans*, except where indicated in the “Genotype” field.

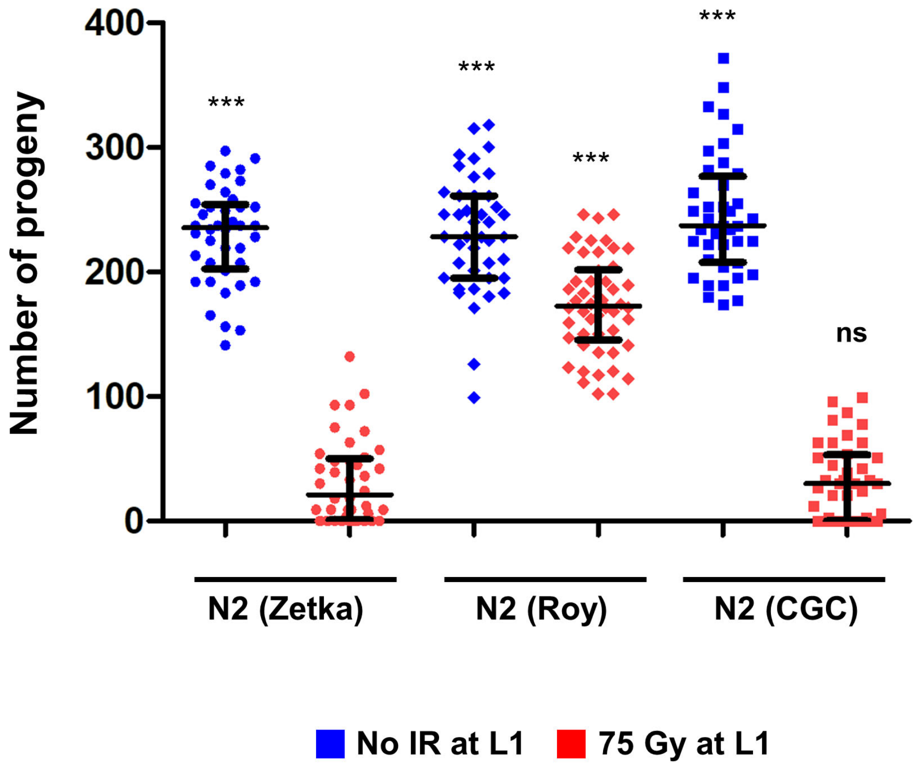
400
300
200
100
0
Number of progeny
***
***
***
***
ns
N2 (Zetka)
N2 (Roy)
N2 (CGC)
No IR at L1
75 Gy at L1

A

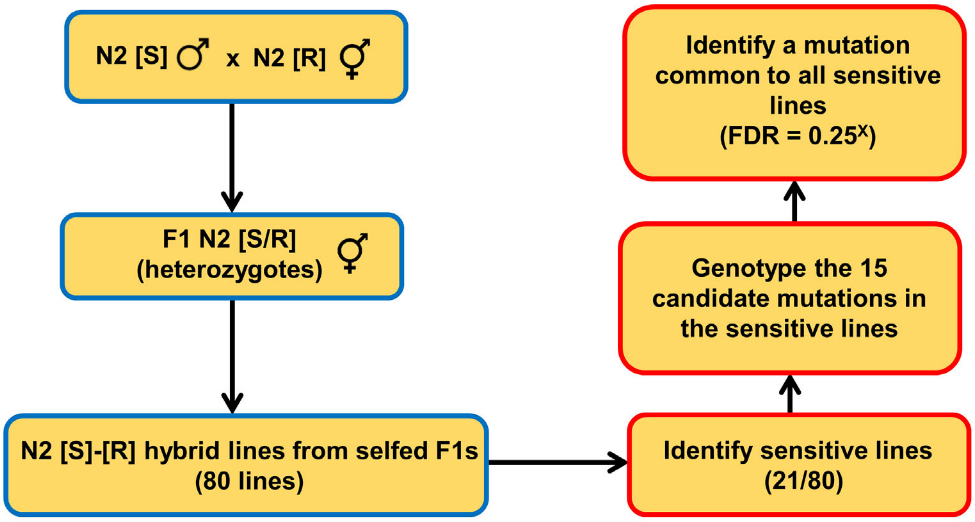
N2 [S] ♂ x N2 [R] ⚥
F1 N2 [S/R] (heterozygotes) ⚥
N2 [S]-[R] hybrid lines from selfed F1s (80 lines)
Identify sensitive lines (21/80)
Genotype the 15 candidate mutations in the sensitive lines
Identify a mutation common to all sensitive lines (FDR = 0.25^X)

B

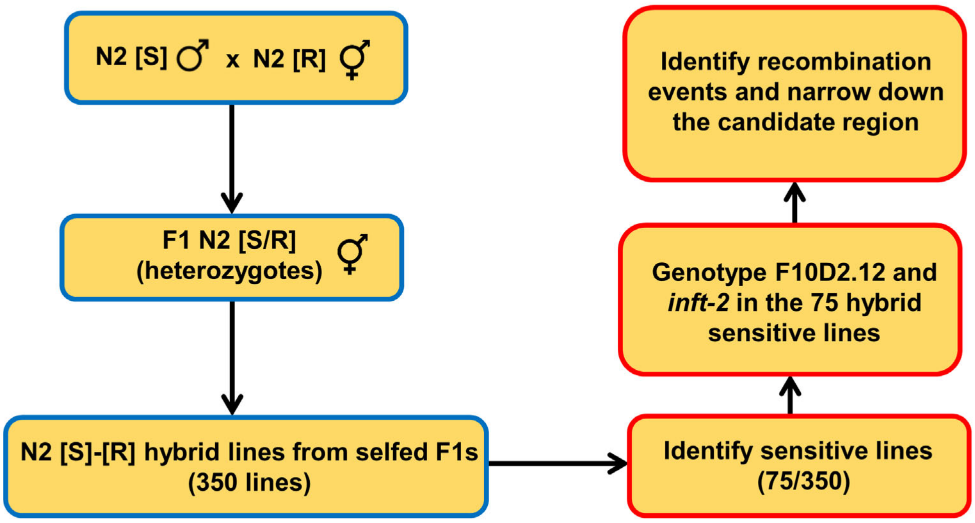
N2 [S] ♂ x N2 [R] ⚥
F1 N2 [S/R] (heterozygotes) ⚥
N2 [S]-[R] hybrid lines from selfed F1s (350 lines)
Identify sensitive lines (75/350)
Genotype F10D2.12 and *inft-2* in the 75 hybrid sensitive lines
Identify recombination events and narrow down the candidate region

C

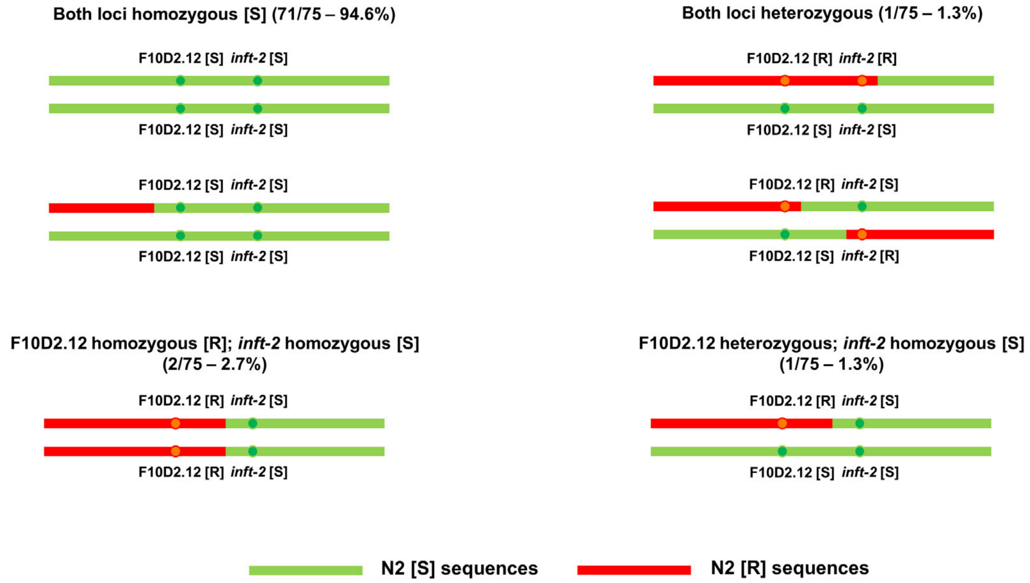
Both loci homozygous [S] (71/75 – 94.6%)
F10D2.12 [S] *inft-2* [S]
F10D2.12 [S] *inft-2* [S]
F10D2.12 [S] *inft-2* [S]
F10D2.12 [S] *inft-2* [S]
Both loci heterozygous (1/75 – 1.3%)
F10D2.12 [R] *inft-2* [R]
F10D2.12 [S] *inft-2* [S]
F10D2.12 [R] *inft-2* [S]
F10D2.12 [S] *inft-2* [R]
F10D2.12 homozygous [R]; *inft-2* homozygous [S] (2/75 – 2.7%)
F10D2.12 [R] *inft-2* [S]
F10D2.12 [R] *inft-2* [S]
F10D2.12 heterozygous; *inft-2* homozygous [S] (1/75 – 1.3%)
F10D2.12 [R] *inft-2* [S]
F10D2.12 [S] *inft-2* [S]
N2 [S] sequences
N2 [R] sequences

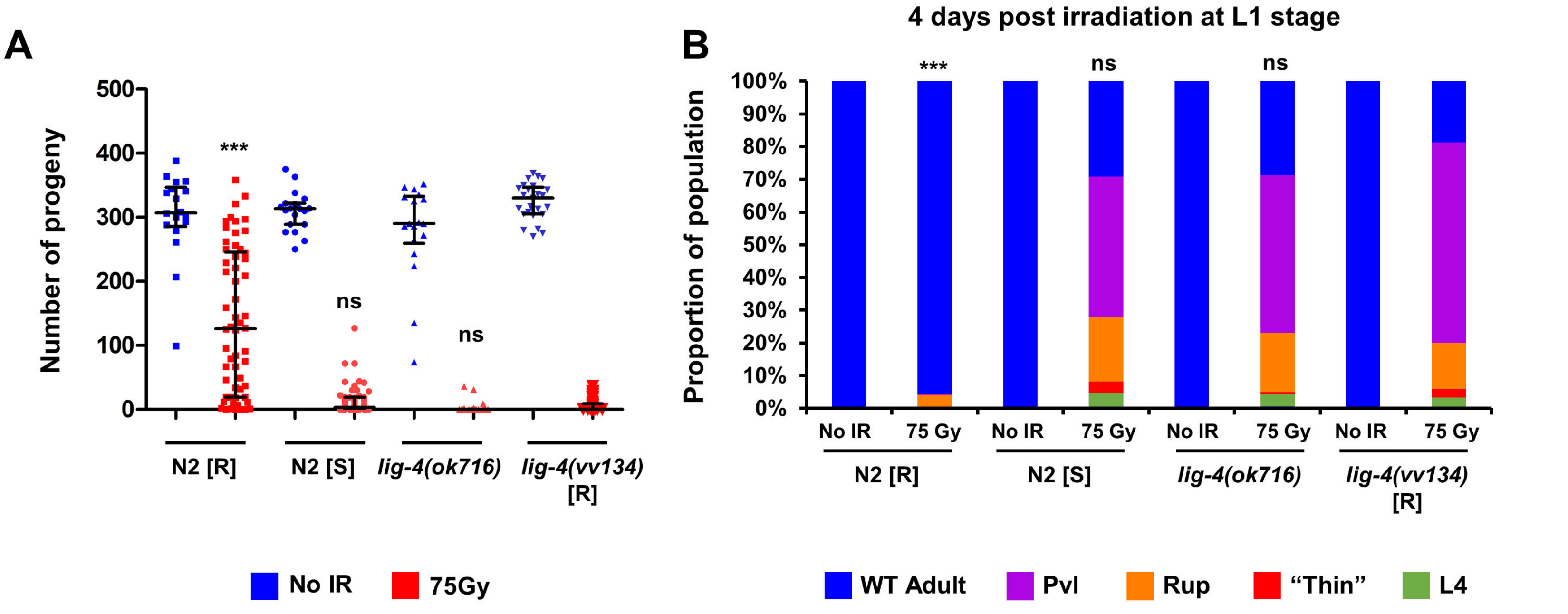
A
Number of progeny
500
400
300
200
100
0
***
ns
ns
N2 [R]
N2 [S]
lig-4(ok716)
lig-4(vv134) [R]
No IR
75Gy
4 days post irradiation at L1 stage
B
Proportion of population
100%
90%
80%
70%
60%
50%
40%
30%
20%
10%
0%
***
ns
ns
No IR
75 Gy
N2 [R]
N2 [S]
lig-4(ok716)
lig-4(vv134) [R]
WT Adult
Pvl
Rup
"Thin"
L4

| **N2 lines** | | |
|---|---|---|
| **Strain name** | **Origin** | **L1 IR response** |
| **N2** | Zetka laboratory | Sensitive |
| **N2** | CGC | Sensitive |
| **N2** | Roy laboratory | Resistant |
| **N2** | Hekimi laboratory | Resistant |
| **N2** | Andersen laboratory | Resistant |
| **N2** | NBRP | Resistant |
| **N2** | Zhen laboratory | Resistant |
| **Other *C. elegans* wild type isolates** | | |
| **Strain name** | **Origin** | **L1 IR response** |
| **QX1211** | San Francisco, California | Resistant |
| **JU775** | Lisbon, Portugal | Resistant |
| **CB4856** | Oahu, Hawaii | Resistant |
| **DL238** | Hawaii, Hawaii | Resistant |
| **MY16** | Mecklenbeck, Germany | Resistant |
| **MY23** | Roxel, Germany | Resistant |
| **EG4724** | Amares, Portugal | Resistant |
| **JU258** | Ribeiro Frio, Madeira | Resistant |
| **LKC34** | Madagascar | Resistant |
| **JU1088** | Kakegawa, Japan | Resistant |
| **ED3073** | Limuru, Kenya | Resistant |
| **DL226** | Corvallis, Oregon | Resistant |
| **AB4** | Adelaide, Australia | Resistant |
| **JU1171** | Concepcion, Chile | Resistant |
| **AB1** | Adelaide, Australia | Resistant |
| **JU1652** | Montevideo, Uruguay | Resistant |
| **JU1896** | Athens, Greece | Resistant |
| ***C. briggsae* strains** | | |
| **Strain name** | **Origin** | **L1 IR response** |
| **VT847** | Hawaii, Hawaii | Resistant |
| **PB800** | Dayton, Ohio | Resistant |

Table S2

| | **Control L1** unhatched/total (%) | **Irradiated L1 (75 Gy)** unhatched/total (%) |
|---|---|---|
| N2 [R] | 7/1111 (0.63%)$^{ns}$ | 55/582 (9.45%)*** |
| N2 [S] | 11/1293 (0.85%) | 4/593 (0.67%) |
| | **Control L4** unhatched/total (%) | **Irradiated L4 (75 Gy)** unhatched/total (%) |
| N2 [R] | 7/922 (0.76%)$^{ns}$ | 617/1139 (54.17%)$^{ns}$ |
| N2 [S] | 6/948 (0.63%) | 665/1297 (51.27%) |

| Gene name | Different from reference in | Mutation | Chromosomal location | Mutation effect | Molecular identity | Potential function |
|---|---|---|---|---|---|---|
| *trpp-8* | N2 [S] | C->A | Chr I, 5616196 | AA substitution | Transport protein | Involved in intracellular transport and meiosis |
| *ZC239.15* | N2 [S] | C->G | Chr II, 3223743 | AA substitution | BTB2 domain containing KCTD10 ortholog | Potassium chanel (inferred) |
| *emb-27* | N2 [S] | G->C | Chr II, 8160254 | AA substitution | APC subunit, cdc16 ortholog | Mitotic Exit |
| *Y22D7AL.7* | N2 [S] | G->A | Chr III, 1578431 | AA substitution | Unknown protein, some homology to Rouxiella DNA binding response regulator | UNKNOWN |
| *C50C3.2* | N2 [S] | G->T | Chr III, 8188649 | AA substitution | SPTAN1 ortholog (Spectrin) | Cell cortex structural |
| *kin-24* | N2 [S] | G->T | Chr IV, 9840056 | AA substitution | Kinase, human FES ortholog | Kinase, SH2 domain containing, PKC superfamily |
| *F07C6.4* | N2 [S] | C->T | Chr IV, 12809030 | AA substitution | FRMD4B ortholog | Putative actin binding, FERM domain |
| *F10D2.12* | N2 [S] | G->A | Chr V, 7149488 | AA substitution | Hexosyl transferase (putative) | Glycosylation (inferred) |
| *inft-2* | N2 [S] | C->A | Chr V, 13003815 | AA substitution | Inverted formin | FH2 superfamily domain, cytoskeletal organization |
| *sru-48* | N2 [S] | C->T | Chr X, 5180366 | AA substitution | Serpentine GPCR receptor | Cell signaling |
| *F48F7.3* | N2 [S] | G->T | Chr X, 13964441 | AA substitution | Galactosyl transferase (putative) | Galactosylation (inferred) |
| *C06G1.6* | N2 [S] | G->A | Chr X, 16652703 | AA substitution | PPP2R3B ortholog (putative phosphatase) | Dephosphorylation (inferred) |
| *fbxb-69* | N2 [S] | T->A | Chr I, 14283531 | STOP gained | F-box protein | UNKNOWN |
| *C33H5.1* | N2 [S] | G->T | Chr IV, 7810427 | STOP gained | Nematode-specific putative methyltransferase | UNKNOWN |
| *bre-5* | N2 [S] | G->T | Chr IV, 12050973 | Splice site loss | Galactosyl transferase (putative) | Galactosylation (inferred) |

| Strain name | Genotype | Source |
|---|---|---|
| **N2 (Zetka)** | Wild type isolate | Obtained from the CGC |
| **N2 (CGC)** | Wild type isolate | Obtained from the CGC |
| **N2 (Roy)** | Wild type isolate | Generously provided by Dr. Richard Roy |
| **N2 (Hekimi)** | Wild type isolate | Generously provided by Dr. Siegfried Hekimi |
| **N2 (Andersen)** | Wild type isolate | Generously provided by Dr. Eric Andersen |
| **N2 (Zhen)** | Wild type isolate | Generously provided by Dr. Mei Zhen |
| **N2 (NBRP)** | Wild type isolate | Obtained from the NBRP of Japan |
| **QX1211** | Wild type isolate | Obtained from the CGC |
| **JU775** | Wild type isolate | Obtained from the CGC |
| **CB4856** | Wild type isolate | Obtained from the CGC |
| **DL238** | Wild type isolate | Obtained from the CGC |
| **MY16** | Wild type isolate | Obtained from the CGC |
| **MY23** | Wild type isolate | Obtained from the CGC |
| **EG4724** | Wild type isolate | Obtained from the CGC |
| **JU258** | Wild type isolate | Obtained from the CGC |
| **LKC34** | Wild type isolate | Obtained from the CGC |
| **JU1088** | Wild type isolate | Obtained from the CGC |
| **ED3073** | Wild type isolate | Obtained from the CGC |
| **DL226** | Wild type isolate | Obtained from the CGC |
| **AB4** | Wild type isolate | Obtained from the CGC |
| **JU1171** | Wild type isolate | Obtained from the CGC |
| **AB1** | Wild type isolate | Obtained from the CGC |
| **JU1652** | Wild type isolate | Obtained from the CGC |
| **JU1896** | Wild type isolate | Obtained from the CGC |
| **VT847** | *C. briggsae* wild type isolate | Obtained from the CGC |
| **PB800** | *C. briggsae* wild type isolate | Obtained from the CGC |
| **NL2098** | *rrf-1(pk1417)* I | Obtained from the CGC |
| **EZ208** | *htp-3(vc75)* I | EMS mutant from TILLING screen |
| **NL2099** | *rrf-3(pk1426)* II | Obtained from the CGC |
| **YY470** | *dcr-1(mg375)* III | Obtained from the CGC |
| **WM158** | *ergo-1(tm1860)* V | Obtained from the CGC |
| **NL2550** | *ppw-1(pk2505)* I | Obtained from the CGC |
| **NL5117** | *ppw-2(pk1673)* I | Obtained from the CGC |
| **WM160** | *sago-1(tm1195)* V | Obtained from the CGC |
| **WM154** | *sago-2(tm894)* I | Obtained from the CGC |
| **MT13649** | *nurf-1(n4295)* II | Obtained from the CGC |
| **RB1831** | *set-22(ok2370)* V | Obtained from the CGC |
| **EU1481** | *orIs20* [*pie-1p::GFP::MOE*] | Generously provided by Dr. Richard Roy |

| Strain name | Genotype | |
|---|---|---|
| **EZ412/N2 [S]** | Wild type | Generated from a single **N2 (CGC)** animal |
| **EZ413/N2 [R]** | Wild type | Generated from a single **N2 (Andersen)** animal |
| **EZ421** | *dcr-1(vv121)* III [S] | Generated by CRISPR mutagenesis in **N2 [S]** |
| **EZ422** | *dcr-1(vv122)* III [R] | Generated by CRISPR mutagenesis in **N2 [R]** |
| **EZ437** | *inft-2(vv135) F10D2.12(vv136)* V [R] | Generated by CRISPR mutagenesis in **N2 [R]** |
| **RB873** | *lig-4(ok716)* III | Obtained from the CGC |
| **EZ436** | *lig-4(vv134)* III [R] | Generated by CRISPR mutagenesis in **N2 [R]** |
| **EZ443** | *lig-4(vv141)* III [S] | Generated by CRISPR mutagenesis in **N2 [S]** |
| **EZ454** | *nhj-1(vv144)* V [R] | Generated by CRISPR mutagenesis in **N2 [R]** |
| **FX01203** | *cku-80(tm1203)* III | Obtained from the NBRP of Japan |
| **HBR1099** | *unc-119(ed3)* III; *goeEx386 [WRM0635D_B04(pRedFlp-Hgr)(H19N07.3[21364]::S0001_pR6K_Amp_2xTY1ce_EGFP_FRT_rpsl_neo_FRT_3xFlag) dFRT::unc-119-Nat]* | Generously provided by Dr. Henrik Bringmann |
| **EZ477** | *unc-119(ed3)* III; *nhj-1(vv144)* V*; goeEx386* | Generated by crossing **HBR1099** with **EZ454** |
| **EZ463** | *cku-80(tm1203)* III; *nhj-1(vv144)* V | Generated by crossing **EZ454** with **FX01203** |
| **EZ459** | *nhj-1(vv147[nhj-1::OLLAS])* V [R] | Generated by CRISPR mutagenesis in **N2 [R]** |
| **EZ461** | *cku-80(tm1203)* III; *nhj-1(vv147[nhj-1::OLLAS])* V | Generated by crossing **YA942** with **EZ459** |
| **EZ460** | *lig-4 (vv134)* III; *nhj-1(vv147[nhj-1::OLLAS])* V | Generated by crossing **EZ436** with **EZ459** |
| **EZ457** | *lig-4(vv145[lig-4::OLLAS])* III [R] | Generated by CRISPR mutagenesis in **N2 [R]** |
| **EZ464** | *cku-80(tm1203) lig-4(vv145[lig-4::OLLAS])* III | Generated by crossing **YA942** with **EZ457** |
| **EZ465** | *lig-4(vv145[lig-4::OLLAS])* III; *nhj-1(vv144)* V | Generated by crossing **EZ454** with **EZ457** |
| **MR156** | *unc-119(ed3)* III*; rrIs01* [*elt-2::GFP unc-119(+)*] X | Generously provided by Dr. Richard Roy |
| **EZ466** | *unc-119(ed3)* III*; nhj-1(vv144)* V; *rrIs01* [*elt-2::GFP unc-119(+)*] X | Generated by crossing **MR156** with **EZ454** |
| **GE4132** | *unc-32(e189) com-1(t1626)/*qC1 [*dpy-19(e1259) glp-1(q339)*] III | Obtained from the CGC |
| **EZ467** | *unc-32(e189) com-1(t1626)/*qC1 [*dpy-19(e1259) glp-1(q339)*] III; *nhj-1(vv144)* V | Generated by crossing **GE4132** with **EZ454** |
| **EZ476** | *unc-32(e189) com-1(t1626) lig-4(vv134)/*qC1 [*dpy-19(e1259) glp-1(q339)*] III | Generated by CRISPR mutagenesis in **GE4132** |
